# A platform for mapping reactive cysteines within the immunopeptidome

**DOI:** 10.1101/2024.04.02.587775

**Authors:** Chenlu Zhang, Chen Zhou, Assa Magassa, Deyu Fang, Xiaoyu Zhang

## Abstract

The major histocompatibility complex class I (MHC-I) antigen presentation pathways play pivotal roles in orchestrating immune responses. Recent studies have begun to utilize cysteines within the immunopeptidome for therapeutic applications, such as using covalent ligands to create haptenated neoantigens for inducing an immune response. In this study, we report a platform for mapping reactive cysteines present on MHC-I-bound peptide antigens. We have developed cell-impermeable sulfonated maleimide probes capable of effectively capturing reactive cysteines on antigens. Utilizing these probes in chemoproteomic experiments, we discovered that cysteines on MHC-I-bound antigens exhibit various degrees of reactivity. Furthermore, interferon-gamma stimulation produces increased reactivity of cysteines at position 8 of 9-mer MHC-I-bound antigens. Our findings may open up new avenues for understanding the distinctive roles of cysteine within the MHC-I immunopeptidome and leveraging the differentially reactive cysteines for therapeutic intervention.

## Introduction

Major histocompatibility complex class I (MHC-I) molecules play a crucial role in presenting peptides derived from cytosolic and nuclear proteins on the cell surface^1^. These peptides, also referred to as antigens, originate from degraded proteins processed by the proteasome^2^. Following degradation, the peptides are translocated into the endoplasmic reticulum (ER), where they bind to MHC-I molecules with the assistance of various chaperone proteins. Human MHC-I molecules are comprised of a membrane-anchored heavy chain encoded by the *HLA-A*, *HLA-B* or *HLA-C* genes, and a light chain called β2-microglobulin. Upon recognition of appropriate peptides, typically consisting of 8-13 amino acids, the peptide-MHC-I complex (pMHC-I) is transported to the cell surface, resulting in the exposure of antigens to the extracellular space^3^. Alternatively, antigens presented by MHC-I can originate from the lysosome-mediated degradation of internalized proteins and microbial pathogens, a process known as cross-presentation that occurs in professional antigen presenting cells (APCs)^4^. The MHC-I antigen presentation pathways play central roles in regulating immune responses, impacting various physiological functions and disease progressions, including autoimmune diseases^5^, viral infections^6^, and cancer immune surveillance^7^. This essential role hinges largely on the interaction between the T-cell receptor (TCR) of cytotoxic CD8^+^ T cells and the pMHC-I complex displayed on the cell surface^8^. In the context of cancer, the identification of tumor-specific or tumor-associated antigens presented on MHC-I is fundamental for mounting an effective CD8^+^ T cell antitumor immune response^9^. Conversely, in autoimmune disorders, autoreactive T cells may evade thymic negative selection and peripheral tolerance mechanisms, leading to recognition of self-derived pMHC-I complexes in otherwise healthy tissues^10^.

Given the fundamental role of pMHC-I complex in physiology, deciphering the antigen sequences presented by MHC-I is imperative for understanding how pMHC-I complexes influence the immune system and for developing novel therapeutic strategies. Computational tools have been extensively employed to predict these antigen sequences^11^. Nevertheless, a limitation of current algorithms is their suboptimal performance when analyzing antigens containing cysteine residues^12^. In human genome, approximately 262,000 cysteines are encoded and distributed within proteins from diverse classes and families^13^. Leveraging the intrinsic reactivity of proteinaceous cysteine, the development of covalent chemical probes and drugs holds immense potential for elucidating protein functions and offering therapeutic interventions^14^. Despite this, our understanding of the functionality and reactivity of cysteines within MHC-I antigens has historically been limited. Recent studies have begun to exploit antigen cysteines for therapeutic applications. For example, introducing a disulfide bond between TCR and MHC-I-bound antigens has been shown to facilitate T cell activation^15^. Additionally, employing covalent inhibitors to modify cysteines on intracellular proteins, such as KRAS-G12C, EGFR-T790M and BTK, leads to the presentation of inhibitor-modified antigens on MHC-I^16, 17^. This unique phenomenon can be therapeutically exploited to recruit cytotoxic T cells to tumor cells presenting these haptenated neoepitopes^16, 17^. Out of the 520,000+ MHC-I-associated antigens predicted in The Immune Epitope Database (IEDB), approximately 20,000 antigens harbor cysteine residues. A fundamental question arises: how many of these cysteines are amenable to therapeutic targeting? However, technologies for systematically mapping and studying cysteines within MHC-I-bound antigens have been lacking. In this study, we have developed a platform for mapping reactive cysteines present on MHC-I-bound antigens. This platform enables the exploration of potentially targetable immunopeptidome from diverse perspectives.

## Results

### Development of cell-impermeable reactivity probes for mapping reactive cysteines within the immunopeptidome

To gain insight into cysteines within the immunopeptidome, we propose developing broad-spectrum probes capable of capturing antigen cysteines for diverse applications. The design of these reactivity probes should incorporate three crucial features: 1) spatial specificity: The probes should remain in the extracellular space, exclusively targeting extracellular cysteines. 2) residue specificity: The probes selectively target cysteine-containing peptide antigens without reacting with other amino acids; and 3) quantification capability: The probes form covalent bonds with antigens, generating unique modifications that can be quantified using various analytical techniques, including flow cytometry, fluorometric assays, and mass spectrometry (MS). It is worth noting that due to the oxidizing environment of the extracellular milieu, resulting in interchain disulfide bond formation among extracellular proteinaceous cysteines^18^, we expect that cell-impermeable cysteine-reactive probes predominantly label cysteines within the immunopeptidome. In accordance with these criteria, we synthesized six probes, each incorporating one of three cysteine-reactive groups (iodoacetamide, chloroacetamide, and maleimide), all linked to desthiobiotin (DTB) (**Fig. 1a**). Among these probes, iodoacetamide-PEG-desthiobiotin (IA-DTB) and desthiobiotin iodoacetamide (DBIA) are widely used for profiling intracellular proteinaceous cysteines using the activity-based protein profiling (ABPP) platform^19, 20^. To potentially achieve spatial specificity, we introduced a negatively charged group, such as carboxyl or sulfonate, into certain probes, a strategy previously used to confer cell impermeability to small molecules^21^.

**Fig. 1.**
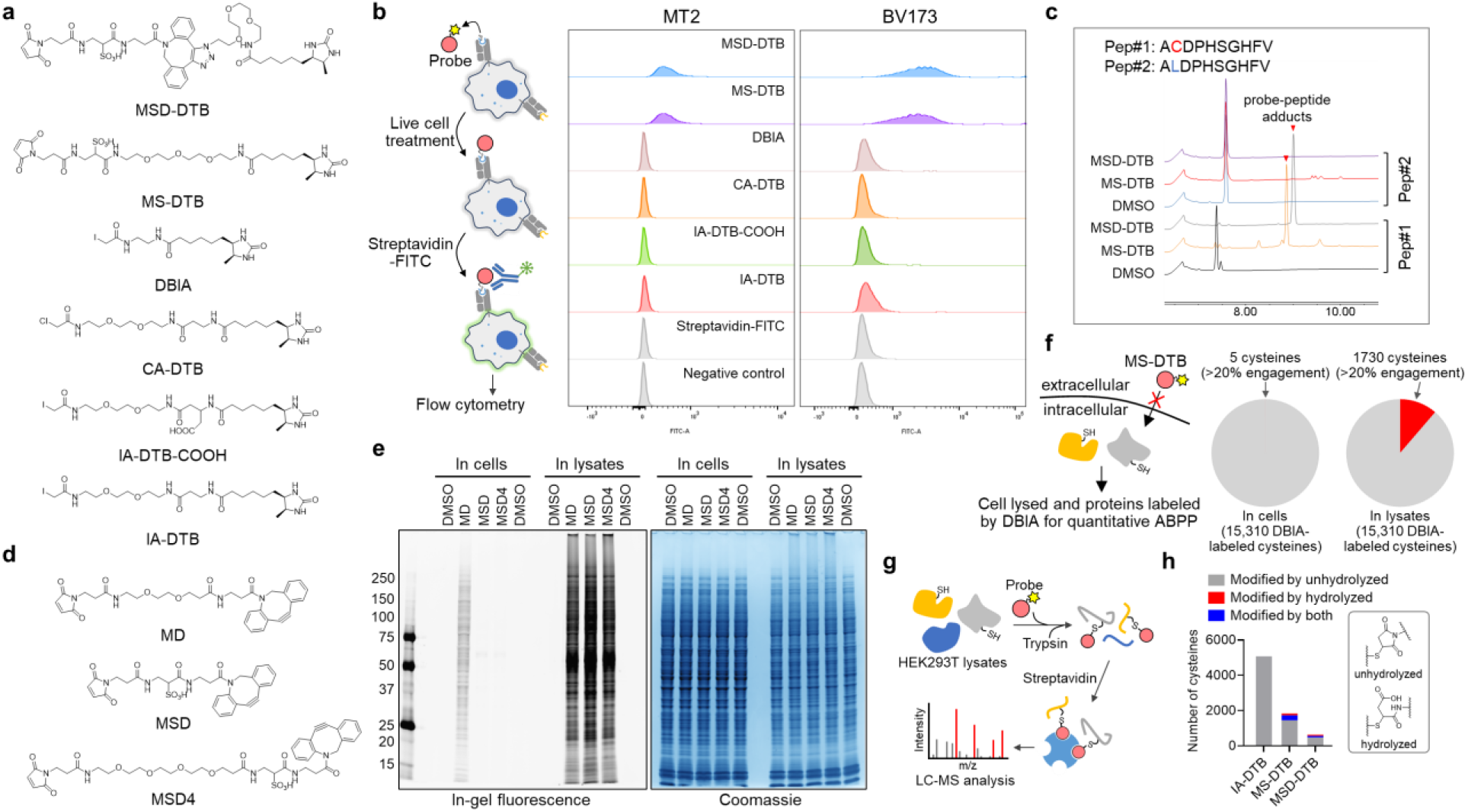
Development of cell-impermeable cysteine-reactive probes. **a**. Structures of 6 reactivity probes, with each one incorporating one of three cysteine-reactive groups (iodoacetamide, chloroacetamide, and maleimide), all linked to desthiobiotin (DTB). **b**. Left panel: schematic representation of using flow cytometry to measure cell surface labeling by the probe. Right panel: MSD-DTB and MS-DTB probes exhibited superior cell surface labeling in BV173 and MT2 cells compared to the others. Cells were treated with 10 µM of each reactivity probe for 30 minutes. The result is a representative of three experiments (n = 3 biologically independent experiments). **c**. LC-MS analysis revealed that MSD-DTB and MS-DTB probes selectively modify cysteine, but not histidine, serine, and N-terminal amine. The result is a representative of two experiments (n = 2 biologically independent experiments). **d**. Structures of MD, MSD, and MSD4, all containing a dibenzocyclooctyne group for the azide-alkyne cycloaddition with a fluorophore. **e**. In-gel fluorescence analyses revealed that sulfonated maleimide probes minimally labeled proteins in live cells, while yielding comparable levels of proteome labeling to its cell-permeable counterpart probe, MD, in cell lysates. The result is a representative of two experiments (n = 2 biologically independent experiments). **f**. Cysteine-directed ABPP analyses revealed that MS-DTB only engaged 5 cysteines with >20% engagement out of 15,310 quantified cysteines in live cells, while it engaged 1730 cysteines with >20% engagement in cell lysates. Cysteine engagement data represent mean values (n = 2 biologically independent experiments). **g**. Schematic representation of employing MSD-DTB and MS-DTB in proteomics workflow to identify probe-modified cysteine-containing peptides. **h**. The number of cysteines modified by IA-DTB, MSD-DTB and MS-DTB. The result is a representative of two experiments (n = 2 biologically independent experiments). Abbreviations: MSD-DTB, maleimide-sulfonate-dibenzocyclooctyne-DTB. MS-DTB, maleimide-sulfonate-DTB. DBIA, desthiobiotin iodoacetamide. IA-DTB, iodoacetamide-PEG-desthiobiotin. CA-DTB, chloroacetamide-PEG-desthiobiotin. IA-DTB-COOH, iodoacetamide-carboxylate-PEG-desthiobiotin.

Initially, we used flow cytometry to assess cell surface labeling by these reactivity probes. BV173 (human B cell leukemia) and MT2 (human T cell leukemia) cells were treated with the probe, followed by washing out of the free probe and subsequent incubation with streptavidin-FITC (**Fig. 1b**, left panel). As streptavidin-FITC remains impermeable to cells^22^, the fluorescence measured by flow cytometry should primarily reflect extracellular cysteine labeling by the probe. The results revealed that two sulfonated maleimide probes, maleimide-sulfonate-dibenzocyclooctyne-DTB (MSD-DTB) and maleimide-sulfonate-DTB (MS-DTB), exhibited superior cell surface labeling in both BV173 and MT2 cells compared to others (**Fig. 1b**, right panel). To validate the specific reactivity towards cysteines by the MSD-DTB and MS-DTB probes, we incubated them with two peptides at physiological pH (7.4): one containing a cysteine and the other lacking it. Liquid chromatography-mass spectrometry (LC-MS) analysis indicated that the maleimide reactive group selectively modifies cysteine residues, displaying no reactivity towards other nucleophilic amino acids, including histidine and serine, and N-terminal amine (**Fig. 1c** and **Fig. S1a**). Moreover, we incubated maleimide-sulfonate-dibenzocyclooctyne (MSD, **Fig. 1d**) with the same two peptides, followed by an azide-alkyne cycloaddition^23, 24^ with tetramethylrhodamine (TAMRA) azide, and observed labeling only on the cysteine-containing peptide (**Fig. S1b**).

To validate cell impermeability of sulfonated maleimide probes, we treated HEK293T cells with MSD and a MSD derivative with a longer linker (maleimide-sulfonate-PEG4-dibenzocyclooctyne, or MSD4), followed by fluorophore conjugation through an azide-alkyne cycloaddition^23, 24^ and in-gel fluorescence analysis. The results indicated minimal proteome labeling compared to a cell-permeable counterpart probe, maleimide dibenzocyclooctyne (MD), that lacks the sulfonate group (**Fig. 1d,e**). Conversely, treating cell lysates with these reactivity probes resulted in comparable levels of proteome labeling (**Fig. 1e**), suggesting that MSD and MSD4 probes are predominantly cell impermeable. Furthermore, we observed that the only uncharged sulfonated maleimide probe, MD, exhibited cytotoxicity compared to the other sulfonated maleimide probes (**Fig. S1c**). This underscores the cell impermeability of sulfonated maleimide probes incapable of accessing a wide range of intracellular proteins, potentially triggering stress and apoptosis^25^. Finally, we employed cysteine-directed ABPP to assess the global cysteine engagement of MS-DTB in cell lysates versus live cells. MS-DTB-treated samples were further labeled by DBIA for the quantification of DBIA-modified cysteine-containing peptides. Reduced enrichment of DBIA-modified cysteine-containing peptides would suggest the potential engagement of MS-DTB on these cysteines. The results indicated that among 15,310 quantified cysteines, 1,730 cysteines showed >20% engagement by MS-DTB in cell lysates, whereas only 5 cysteines showed >20% engagement by MS-DTB in live cells (**Fig. 1f** and **Supplementary Table 1**). This suggests that MS-DTB has restricted access to intracellular proteins. Collectively, these findings indicate that sulfonated maleimide probes are predominantly cell impermeable.

Next, we sought to investigate the compatibility of sulfonated maleimide probes with chemical proteomics workflow, specifically evaluating whether the probe-modified peptides can be effectively ionized and identified by orbitrap and ion trap mass analyzers. We incubated HEK293T cell lysates with MS-DTB and MSD-DTB probes, followed by trypsin digestion, enrichment with Streptavidin agarose beads, and subsequent analysis on a Tribrid mass spectrometer (**Fig. 1g**). Our comparative analysis demonstrated the effective identification of cysteines modified by both sulfonated maleimide probes (**Fig. 1h** and **Supplementary Table 2**), with MS-DTB probe exhibiting greater coverage than MSD-DTB probe (1833 versus 629 probe-modified cysteine-containing peptides). This difference is likely due to the enhanced ionization of MS-DTB (molecular weight 690.8) compared to MSD-DTB (molecular weight 993.1). Despite carrying a negatively charged sulfonate moiety, we observed a normal isotopic envelope profile of probe-modified peptides in positive mode on the orbitrap mass analyzer (**Fig. S1d**). It is recognized that maleimide-modified cysteines can exist in both unhydrolyzed and hydrolyzed forms^26^. Our data revealed that approximately 20% of probe-modified peptides bear hydrolyzed maleimide (**Fig. 1h**). Therefore, integrating both forms into the proteomics analysis pipeline would be a comprehensive approach for future proteomics studies.

### Sulfonated maleimide probes modify cysteines present on MHC-I-bound antigens

Despite the development of various approaches for mapping peptide antigens^27^, identifying cysteine-containing antigens is known to be challenging^28^. We selected three cell lines – BV173, MT2 and MDA-MB-231 – to examine the potential of sulfonated maleimide probes in capturing cysteine-containing MHC-I bound antigens, thereby facilitating investigations of these enigmatic components. These three cell lines harbor commonly occurring HLA alleles found in worldwide populations (BV173: HLA-A*02:01,30:01; MDA-MB-231: HLA-A*02:01,02:17; and MT2: HLA-A*24:02)^29, 30^. Additionally, according to the TRON Cell Line Portal and Cancer Cell Line Encyclopedia (CCLE), BV173 and MDA-MB-231 cells exhibit high expression levels of HLA genes and MHC-I proteins^30, 31^. Our proteomics and flow cytometry studies indicated that MT2 cells also exhibited high MHC-I expression levels (see below). Thus, these cell lines serve as ideal models for studying potentially abundant cysteines within the MHC-I-associated immunopeptidome. To create control cell lines for parallel comparison, we used CRISPR-Cas9 to knockout all six endogenous *HLA-A*, *HLA-B*, and *HLA-C* genes in these cell lines (referred to as *HLA* knockout hereafter). The knockout of *HLA* genes and the disruption of MHC-I proteins were confirmed through quantitative global proteomics and flow cytometry analysis (**Fig. 2a**, **Fig. S2a,b**, and **Supplementary Table 3**).

**Fig. 2.**
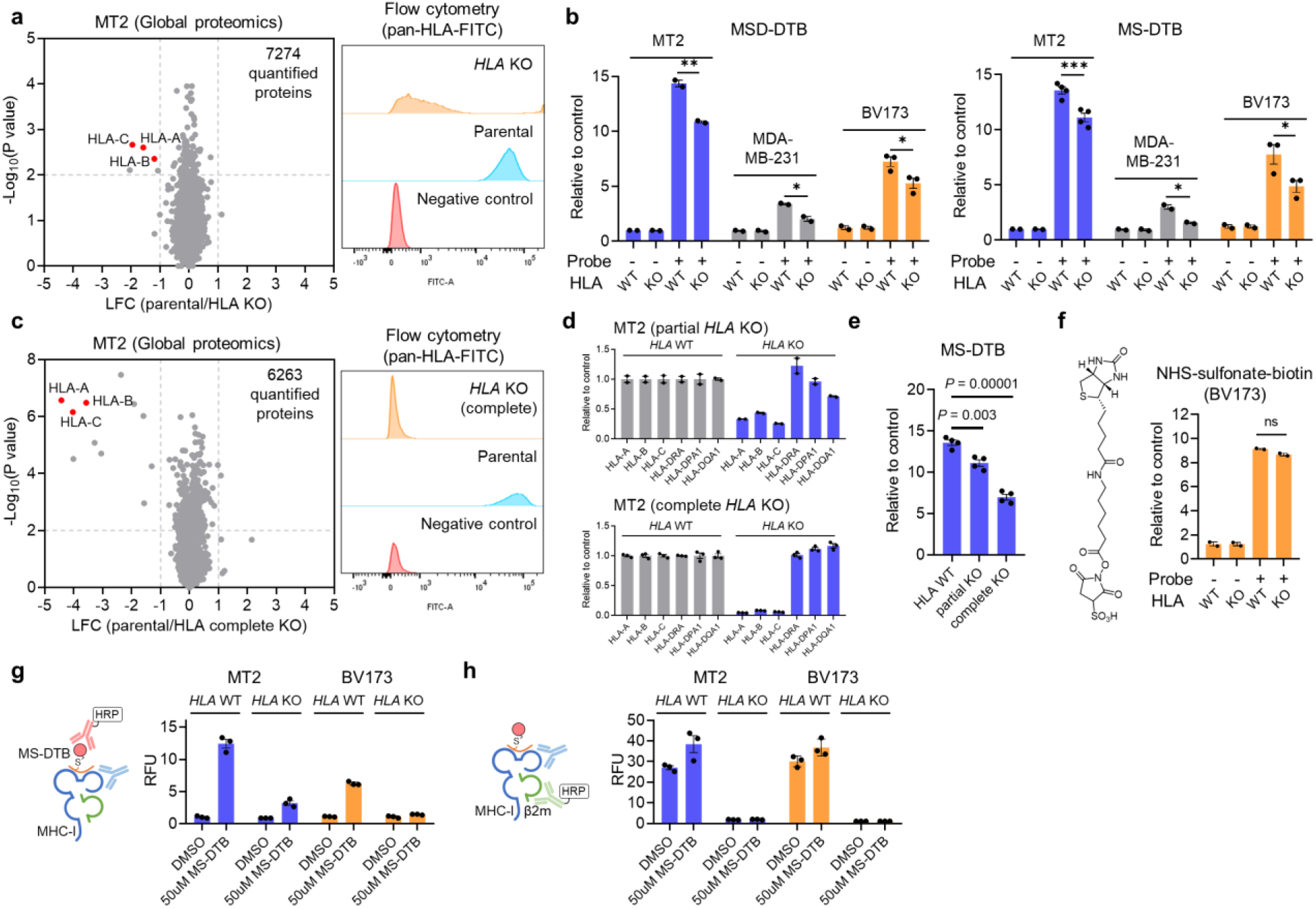
Sulfonated maleimide probes modify cysteines present on MHC-I-bound antigens. **a**. Quantitative global proteomics and flow cytometry studies confirmed the knockout of *HLA-A*, *HLA-B*, and *HLA-C* in MT2 cells (n = 2 biological independent samples for global proteomics, n = 3 biological independent samples for flow cytometry). **b**. Flow cytometry analyses measuring Streptavidin-FITC staining on cells treated with the MSD-DTB or MS-DTB probe (50 µM, 30 minutes). Data represents mean values ± SEM (n = 2 biological independent samples for flow cytometry analysis in MT2 and MDA-MB-231 cells treated with MSD-DTB, n = 2 biological independent samples for flow cytometry analysis in MDA-MB-231 cells treated with MS-DTB, n = 3 biological independent samples for flow cytometry analysis in BV173 cells treated with MSD-DTB and MS-DTB, n = 4 biological independent samples for flow cytometry analysis in MT2 cells treated with MS-DTB). **c**. Quantitative global proteomics and flow cytometry studies confirmed the complete knockout of *HLA-A*, *HLA-B*, and *HLA-C* in MT2 cells (n = 3 biological independent samples for global proteomics, n = 3 biological independent samples for flow cytometry). **d**. Bar graph quantification from global proteomics indicates the knockout of *HLA-A*, *HLA-B*, *HLA-C*, but not genes encoding for MHC-II (*HLA-DRA*, *HLA-DPA1*, *HLA-DQA1*). Data represents mean values ± SEM (n = 2 biological independent samples for global proteomics in Fig. 2a, n = 3 biological independent samples for global proteomics in Fig. 2c). **e**. Flow cytometry analyses measuring Streptavidin-FITC staining on MT2 parental, partial *HLA* knockout, and complete *HLA* knockout cells treated with MS-DTB (50 µM, 30 minutes). Data represents mean values ± SEM (n = 4 biological independent samples for flow cytometry analysis). **f**. Structure of sulfonated NHS-biotin and flow cytometry analyses measuring Streptavidin-FITC staining on BV173 parental and *HLA* knockout cells treated with the sulfonated NHS-biotin probe (50 µM, 30 minutes). Data represents mean values ± SEM (n = 2 biological independent samples for flow cytometry analysis). **g**. ELISA assay measuring the proximity between MHC-I and desthiobiotin with or without MS-DTB treatment in MT2 and BV173 cells. Data represents mean values ± SEM (n = 3 biological independent samples for flow cytometry analysis). **h**. ELISA assay measuring MHC-I and β2-microglobulin assembly with or without MS-DTB treatment in MT2 and BV173 cells. Data represents mean values ± SEM (n = 3 biological independent samples for flow cytometry analysis).

We employed two complementary approaches to assess sulfonated maleimide probes interacting with cysteines on MHC-I-bound antigens. In the first approach, we compared cell surface labeling by the probe in wildtype versus *HLA* knockout cells via flow cytometry. The results revealed a significant decrease in cell surface labeling by both MSD-DTB and MS-DTB probes in *HLA* knockout cells compared to wildtype cells across all three cell lines (**Fig. 2b**), suggesting that the reduced cell surface labeling is likely attributable to MHC-I-bound antigens. We noticed a partial *HLA* knockout in MT2 cells, as indicated by global proteomics and flow cytometry analyses (**Fig. 2a**). While the remaining MHC-I can still contribute to cell surface labeling by the probe, it allows us to further validate the interaction between the probe and antigen cysteines. To this end, we sorted MHC-I negative *HLA* partial knockout cells and generated *HLA* complete knockout cells (**Fig. 2c,d** and **Supplementary Table 3**). Subsequently, we observed a further decrease in cell surface labeling by MS-DTB in *HLA* complete knockout cells (**Fig. 2e**), suggesting that a substantial fraction of the cell surface labeling by the MS-DTB probe is attributed to MHC-I-bound antigens. Furthermore, we measured cell surface labeling by a sulfonated lysine-reactive probe, N-hydroxysuccinimide (NHS)-sulfonate-biotin, and observed similar cell surface labeling in BV173 wildtype and *HLA* knockout cells (**Fig. 2f** and **Fig. S2c**). Given that the majority of lysines in the extracellular milieu are from proteinaceous lysines, the cell surface labeling by the lysine-reactive probe is likely predominantly attributed to protein lysine residues and minimally affected by *HLA* knockout. Therefore, this result serves as an additional critical control, further indicating that the cell surface labeling by the sulfonated maleimide probes is from MHC-I-bound antigens. We noticed that in BV173 and MT2 *HLA* knockout cells, the MSD-DTB and MS-DTB probes still generated cell surface labeling compared to the no probe control, while in MDA-MB-231 cells, this background signal is minimal (**Fig. 2b,e**). One potential explanation is that the sulfonated maleimide probes could interact with MHC-II-bound antigens, leading to labeling in *HLA* knockout cells. The CCLE proteomics data, which includes BV173 and MDA-MB-231 cells but not MT2 cells, indicate that multiple MHC-II proteins in BV173 cells exhibit high expression compared to those in MDA-MB-231 cells (**Fig. S2d**)^31^. This discrepancy may explain the higher background cell surface labeling signal in BV173 cells than in MDA-MB-231 cells.

In the second approach, we employed an enzyme-linked immunosorbent assay (ELISA) to quantify probe-modified antigens within the pMHC-I complex, specifically assessing the proximity between MHC-I and desthiobiotin from the sulfonated maleimide probes. Given our demonstration that sulfonated maleimide probes do not penetrate cells and both MHC-I and β2-microglobulin lack unmodified cysteines in their extracellular domains, the observed proximity between MHC-I and desthiobiotin would imply the interaction of the probe with cysteines on MHC-I-bound antigens. We treated MT2 and BV173 wildtype and *HLA* knockout cells with the MS-DTB probe, lysed the cells, enriched MHC-I protein using a pan-MHC antibody conjugated on the plate, and detected the abundance of desthiobiotin using Streptavidin-HRP (**Fig. 2g**). The results revealed the proximity between MHC-I and desthiobiotin in the presence of MS-DTB in wildtype, but not *HLA* knockout cells (**Fig. 2g**). Moreover, the MS-DTB treatment did not affect pMHC-I folding, as indicated by the ELISA assay measuring the interaction between MHC-I and β2-microglobulin (**Fig. 2h**). Collectively, the findings from both approaches suggest that sulfonated maleimide probes can modify cysteines on MHC-I-bound antigens.

### MHC-I immunopeptidome exhibits distinct cysteine reactivities

Immunopeptidomics has greatly advanced our fundamental understanding of antigenic repertoires associated with various physiological processes and disease states^32^. To gain deeper insight into antigen cysteines modified by sulfonated maleimide probes, we employed immunopeptidomics to examine the abundance and positioning of probe modified cysteines within the MHC-I immunopeptidome. For this investigation, we focused on the use of MS-DTB probe, which yielded a greater number of probe-modified peptides in proteomics studies compared to the MSD-DTB probe (**Fig. 1h**). MT2 and BV173 cells were treated with MS-DTB, followed by cell lysis and immunoprecipitation using a pan-MHC-I antibody to enrich the pMHC-I complex. The immunopeptidome was eluted by trifluoroacetic acid and subsequently analyzed via MS (**Fig. 3a**). During the data search, we included two dynamic modifications on cysteines: 1) MS-DTB (both unhydrolyzed and hydrolyzed forms); and 2) cysteinylation, a frequently observed cysteine modification in immunopeptidomics studies^33^. We identified 3,311 and 2,080 8-13-mer peptides in MT2 and BV173 cells, respectively (**Fig. 3b** and **Supplementary Table 4**). Parallel immunopeptidomics conducted in *HLA* knockout cells resulted in no 8-13-mer peptides identified in both cell lines (**Fig. S3a,b** and **Supplementary Table 4**). In addition, motif analysis of 9-mer peptides, the most preferred length for MHC-I^34^ (1,928 out of 3,331 antigens in MT2 cells, and 1,321 out of 2,080 antigens in BV173 cells, **Fig. S3c**), revealed a close alignment with reported distribution patterns associated with the corresponding *HLA* alleles (**Fig. 3c,d**)^35^, indicating the effective enrichment of the MHC-I immunopeptidome.

**Fig. 3.**
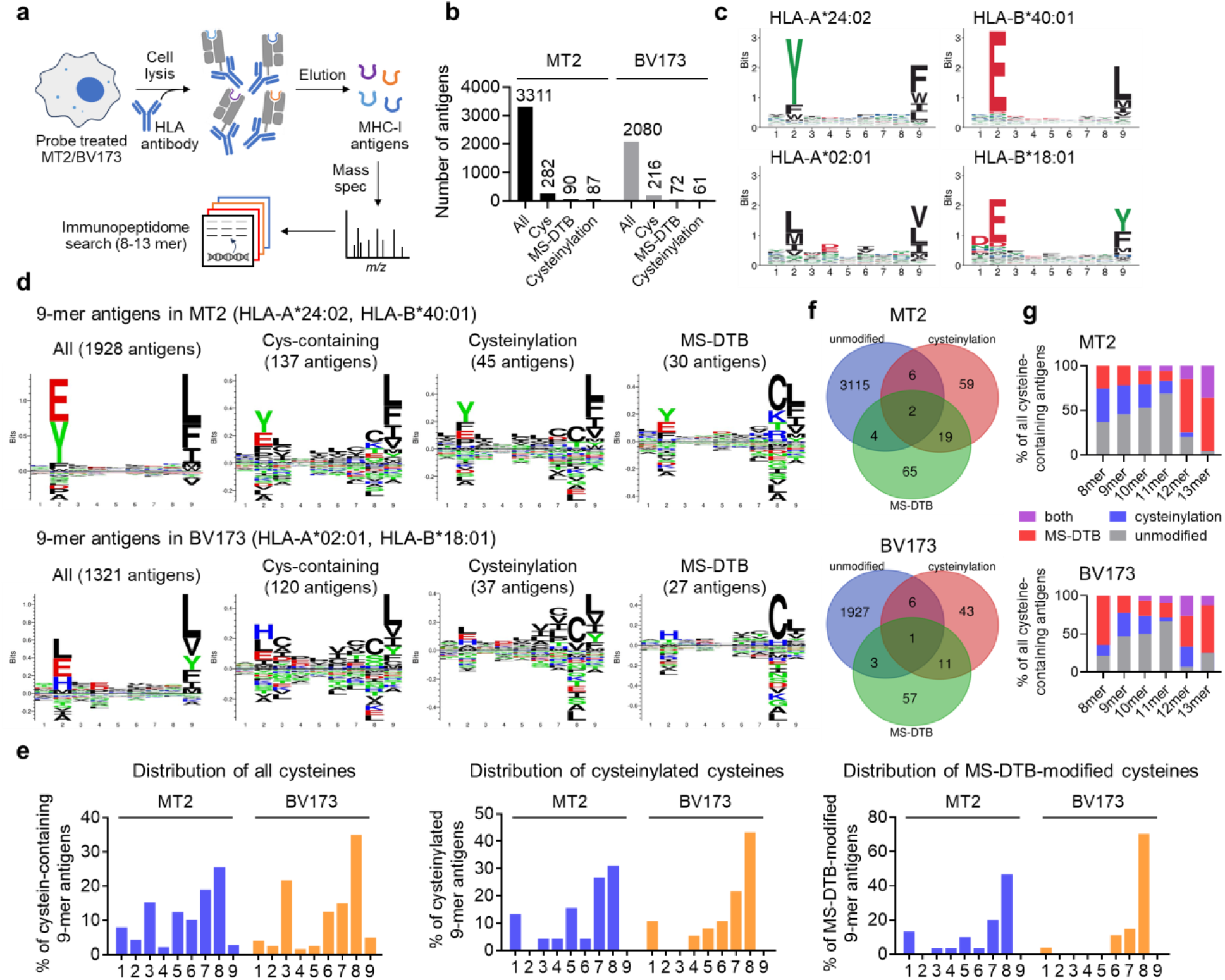
MHC-I immunopeptidome exhibits distinct cysteine reactivities. **a**. Schematic representation of immunopeptidomics. **b**. Number of 8-13-mer MHC-I-associated antigens in MT2 and BV173 cells. The result is a representative of two experiments (n = 2 biologically independent experiments). **c**. Distribution motif of 9-mer antigens associated with HLA-A*24:02, HLA-B*40:01 (MT2) and HLA-A*02:01, HLA-B*18:01 (BV173) (http://mhcmotifatlas.org/home). **d**. Motif analysis of all 9-mer MHC-I-bound antigens, cysteine-containing 9-mer MHC-I-bound antigens, and cysteinylated or MS-DTB-modified 9-mer MHC-I-bound antigens. The result is a representative of two experiments (n = 2 biologically independent experiments). **e**. Distribution of all cysteines, cysteinylated cysteines, and MS-DTB-modified cysteines on 9-mer antigens in MT2 and BV173 cells. The result is a representative of two experiments (n = 2 biologically independent experiments). **f**. Venn diagram of MHC-I-bound antigens with unmodified, cysteinylated, and MS-DTB-modified cysteines. The result is a representative of two experiments (n = 2 biologically independent experiments). **g**. Percentage of unmodified, cysteinylated and MS-DTB-modified 8-13-mer antigens.

Among these antigens, 282 and 216 contain cysteine residues in MT2 and BV173 cells, respectively (**Fig. 3b**). Motif analysis of cysteine-containing 9-mer antigens revealed a similar pattern to that of all 9-mer antigens (**Fig. 3d**, second panel from the left), indicating the confident identification of these cysteines on MHC-I-bound antigens. The minimal enrichment of cysteines at positions 2 and 9 also aligns with the uniquely necessary amino acids at these anchoring positions (**Fig. 3c**), which play a key role in the effective recognition by MHC-I. Intriguingly, other than the anchoring positions 2 and 9, cysteines are not evenly distributed (**Fig. 3e**, first panel from the left). Additionally, the two cell lines harboring different HLA alleles exhibit slightly different distribution patterns of cysteines on antigens. In MT2 cells, cysteines are preferentially located at positions 3, 5, 6, 7 and 8, while in BV173 cells, cysteines are predominantly present at positions 3, 6, 7, and 8 (**Fig. 3e**). Our analysis of cysteinylation on MHC-I-bound antigens revealed enrichment primarily at positions 7 and 8 (**Fig. 3d**, third panel from the left), consistent with the distribution motif reported in a recent study^33^. Approximately 30% of the identified cysteines in both cell lines were found to be modified by MS-DTB (**Fig. 3b**). Notably, there was minimal overlap between MS-DTB-modified and unmodified cysteine-containing antigens (**Fig. 3f**), suggesting that a portion of cysteine-containing antigens may harbor unreactive cysteines that are inaccessible to even highly reactive and high concentration (50 µM) of maleimide probes. Conversely, the cysteines modified by MS-DTB may exist in a solvent-exposed reactive state, making them effectively trackable by maleimide probes. Consistent with this hypothesis, among all the identified cysteines, there is a preference for MS-DTB labeling at positions 1, 5, 7 and 8 in MT2 cells, and positions 6, 7 and 8 in BV173 cells (**Fig. 3e**), indicating the differential presence of reactive cysteines in distinct biological contexts, potentially caused by different MHC-I molecules encoded by various *HLA* alleles. Moreover, it is established that longer peptides (12- and 13-mers) binding to MHC-I molecules occur less frequently than 9-mer antigens^36^. Our immunopeptidomics studies support this observation (**Fig. S3c**). Analysis of cysteine-containing longer antigens revealed that a majority of them were modified by MS-DTB (**Fig. 3g** and **Fig. S3d**). We speculate that these longer cysteine-containing antigens may lack stable associations with MHC-I molecules, resulting in rapid turnover of the pMHC-I complex. However, sulfonated maleimide probes may stabilize them by modifying the cysteines, thereby allowing for more frequent identification of probe-modified longer antigens.

### Using sulfonated maleimide probes to assess alternations in reactive cysteines on MHC-I-bound antigens

MHC-I antigen presentation undergoes tight regulation through various mechanisms occurring during both transcriptional and post-translational stages^37^. Additionally, the presented antigens can be influenced by the induction of immunoproteasomes, potentially generating distinct sequences recognizable by MHC-I molecules^38^. To explore potential alterations in reactive cysteines on MHC-I-bound antigens, we treated BV173 cells with interferon-gamma (IFNγ), a cytokine known to stimulate immunoproteasome generation, and investigated reactive cysteines on MHC-I-bound antigens (**Fig. 4a**). We confirmed IFNγ-induced immunoproteasome generation by measuring increased PSMB9 expression, a subunit replacing PSMB6 upon IFNγ stimulation (**Fig. 4b**)^38^. We also noted a moderate increase in MHC-I expression, consistent with previous findings^39^. Subsequently, using ELISA assays, we observed significantly enhanced MS-DTB engagement within the pMHC-I complex in *HLA* wildtype, but not knockout cells (**Fig. 4c**). This increased engagement could result from increased cysteine reactivity or abundance, especially considering the elevated MHC-I expression, which leads to the presentation of more antigens. Further investigation via comparative immunopeptidomics revealed 2,530 8-13-mer antigens in IFNγ-stimulated cells versus 2,002 in non-stimulated BV173 cells (**Fig. 4d** and **Supplementary Table 5**), indicating an overall enhancement of MHC-I presented antigens with IFNγ stimulation. Consequently, the number of MS-DTB-modified 8-13-mer antigens also increased upon IFNγ simulation (**Fig. 4d**). Motif analysis of 9-mer antigens comparing IFNγ-stimulated versus non-stimulated cells showed a consistent pattern (**Fig. 4e**), suggesting that IFNγ stimulation does not alter the overall pattern of MHC-I presented antigens. Interestingly, IFNγ stimulation increased MS-DTB labeling primarily at position 8 of 9-mer antigens (**Fig. 4e-g**), while other modified positions remained unaffected, further supporting the higher reactivity of cysteines at position 8 within the context of 9-mer antigens associated with *HLA-A2*-encoded MHC-I. Collectively, these findings demonstrate the use of sulfonated maleimide probes in mapping changes in reactive cysteines on MHC-I-bound antigens during altered physiological processes.

**Fig. 4.**
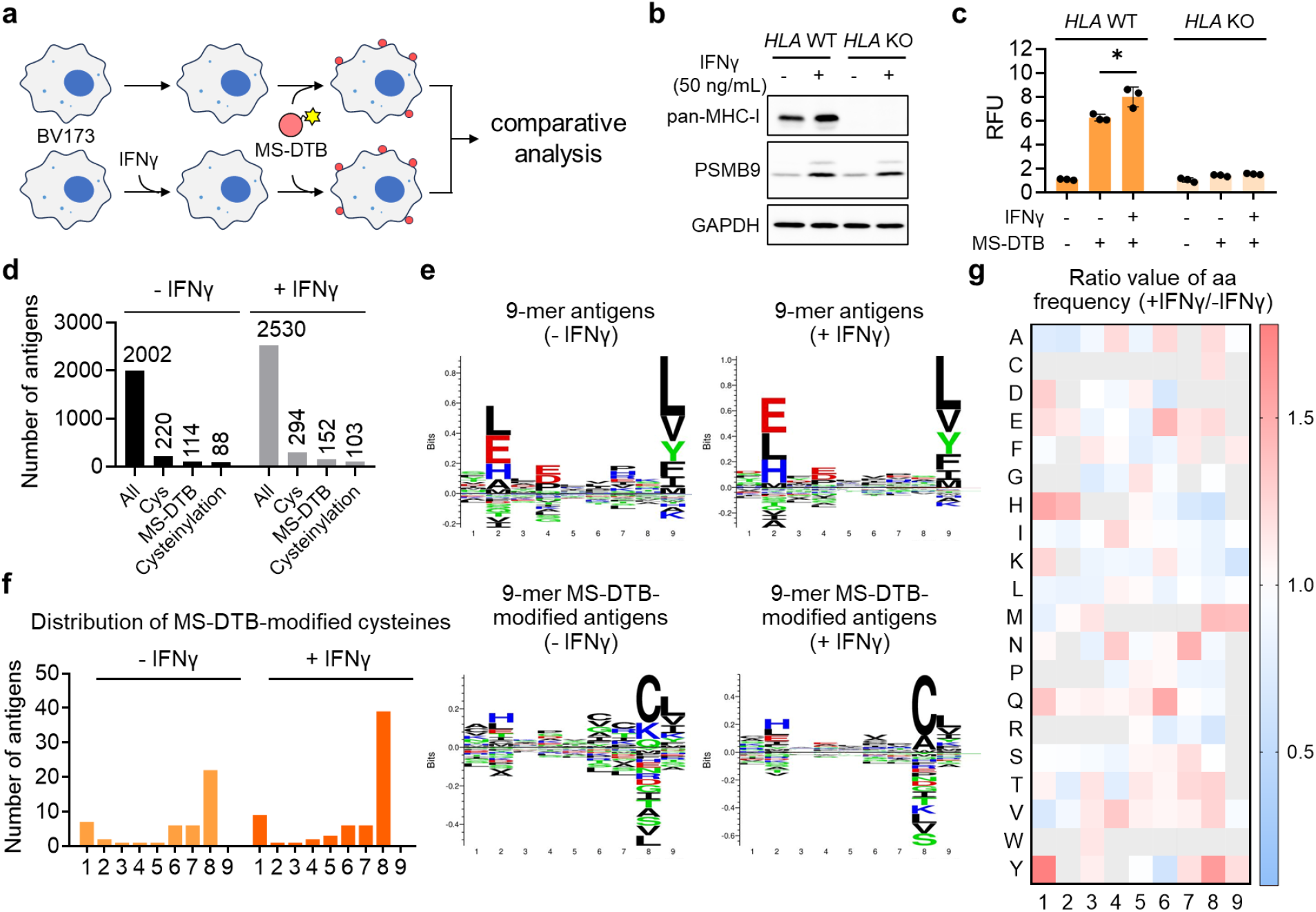
MHC-I immunopeptidome exhibits distinct cysteine reactivities. **a**. Schematic representation of comparative analysis in IFNγ-stimulated versus non-stimulated BV173 cells. **b**. Western blot analysis of MHC-I and PSMB9 expression in BV173 cells stimulated with IFNγ (50 ng/mL, 24 hours). The result is a representative of two experiments (n = 2 biologically independent experiments). **c**. ELISA assay measuring the proximity between MHC-I and desthiobiotin with or without IFNγ stimulation in BV173 cells. Data represents mean values ± SEM (n = 3 biological independent samples for flow cytometry analysis). **d**. Number of 8-13-mer MHC-I-associated antigens in IFNγ-stimulated versus non-stimulated BV173 cells. The result is a representative of two experiments (n = 2 biologically independent experiments). **e**. Motif analysis of all 9-mer MHC-I-bound antigens and MS-DTB-modified 9-mer MHC-I-bound antigens. The result is a representative of two experiments (n = 2 biologically independent experiments). **f**. Distribution of MS-DTB-modified cysteines on 9-mer antigens in IFNγ-stimulated versus non-stimulated BV173 cells. The result is a representative of two experiments (n = 2 biologically independent experiments). **g**. Heatmap showing the ratio values of amino acid (aa) frequency in IFNγ-stimulated versus non-stimulated cells. The percentage of each aa relative to all amino acids at each position was calculated to derive the ratio values between IFNγ-stimulated and non-stimulated cells. A gray shade indicates aa with a percentage below 2%.

### Mapping reactive cysteines on MHC-I-bound antigens using sulfonated maleimide probes via the single-chain trimer model

Studying the immunopeptidome often faces limitations due to the low copy number of pMHC-I complexes on the cell surface^37^, posing challenges for techniques such as proteomics that typically require a substantial number of cells (> 10^8^ cells)^40, 41^. This hurdle becomes especially prominent when investigating individual MHC-I-bound antigens. An alternative method for investigating pMHC-I complexes involves employing single-chain trimer (SCT) models. These engineered constructs consist of covalently linked single chains of MHC-I, β2-microglobulin, and displayed antigenic peptides (**Fig. 5a**)^42^. SCTs maintain structural integrity comparable to native pMHC-I complexes and are capable of robustly activating cytotoxic T cells^43, 44^, making them valuable models for studying virus infection and tumor immunity^43, 44^. Importantly, SCTs can be readily designed to investigate either individual antigens or diverse antigen mixtures within a library^45^. In our study, employing SCTs can facilitate the precise generation of constructs encoding pMHC-I presenting specific cysteine-containing antigens. The proteins encoded by these constructs can be modified using sulfonated maleimide probes for various applications, such as enrichment and in-gel visualization. To illustrate this, we created two SCTs of HLA-A*02:01 presenting previously reported KRAS neoantigens: one containing the G12C mutation (KLVVVGACGV)^16, 17^ and the other containing the G12D mutation (KLVVVGADGV)^46^ (**Fig. 5b**). Both SCT constructs were incorporated with a C-terminal intracellular FLAG tag for enrichment. HEK293T cells were transfected with KRAS-G12C- or KRAS-G12D-SCT and treated with the MSD probe. After cell lysis, SCTs were enriched via FLAG immunoprecipitation, followed by an azide-alkyne cycloaddition^23, 24^ with tetramethylrhodamine (TAMRA) azide, enabling visualization of MSD-modified SCTs via in-gel fluorescence. The results revealed specific labeling of KRAS-G12C-SCT by the MSD probe, while KRAS-G12D-SCT showed no detectable labeling (**Fig. 5c**). Since there are no extracellular unmodified cysteines on MHC-I, β2-microglobulin, and linker components in the SCT constructs (**Fig. S4**), and we have demonstrated that MSD was impermeable to cells (**Fig. 1e**), the labeling signal is likely attributed to the acquired cysteine within the KRAS-G12C neoantigen. Furthermore, we conducted an ELISA assay by incubating sulfonated maleimide probes with recombinant pMHC-I (*HLA-A2*) bound to the KRAS-G12C neoantigen (KLVVVGACGV). The data indicate that probe incubation did not affect the proper folding of the pMHC-I complex involving the KRAS-G12C neoantigen (**Fig. 5d**). Modeling studies using HLA-A*02:01 revealed that Y7 and Y171 on MHC-I formed hydrogen bonds with the N-terminal NH of KRAS-G12C neoantigen. On the other end, D77, Y84 and T143 in the F-pocket of MHC-I formed hydrogen bonds with the C-terminal valine (**Fig. 5e**). The overall confirmation of the KRAS-G12C neoantigen led to the exposure of cysteine at position 8 to the solvent, suggesting the potential for its reactive state to be accessed by the sulfonated maleimide probe.

**Fig. 5.**
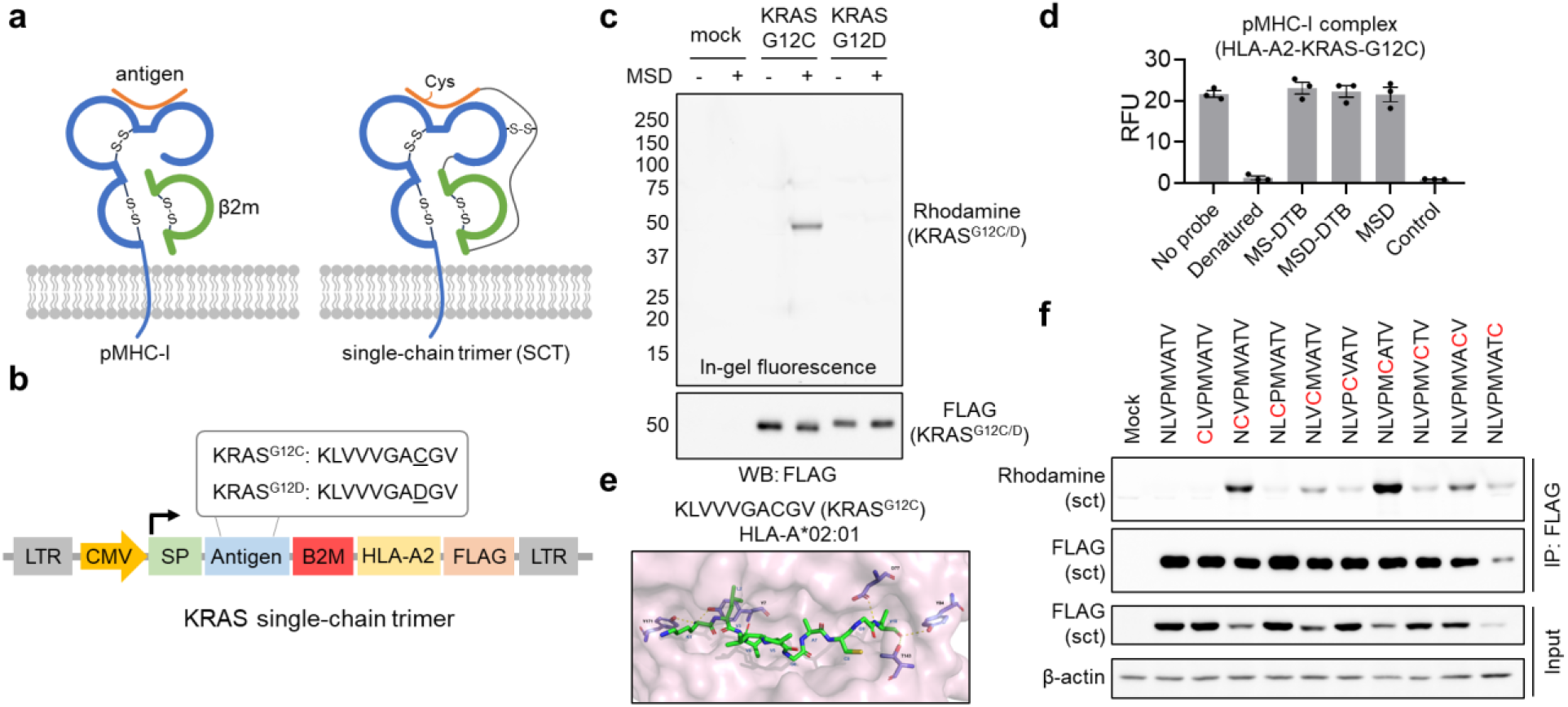
Mapping reactive cysteines on MHC-I-bound antigens using sulfonated maleimide probes via the single-chain trimer model. **a**. Schematic representation of native pMHC-I complex and single-chain trimer (SCT). **b**. Constructs of SCTs of HLA-A*02:01 presenting KRAS-G12C or KRAS-G12D neoantigens. **c**. In-gel fluorescence analysis of KRAS-G12C-SCT and KRAS-G12D-SCT from HEK293T cells treated with the MSD probe (50 µM, 30 minutes). The result is a representative of three experiments (n = 3 biologically independent experiments). **d**. ELISA assay measuring MHC-I and β2-microglobulin assembly in the presence of various sulfonated maleimide probes. Data represents mean values ± SEM (n = 3 biologically independent experiments). **e**. Modeling studies suggest that the cysteine residue within the KRAS^G12C^ neoantigen, when bound to MHC-I, is exposed to the solvent. **f**. In-gel fluorescence analysis of SCTs of HLA-A*02:01 presenting pp65 antigens with cysteines introduced at positions 1-9 from HEK293T cells treated with the MSD probe (50 µM, 30 minutes). The result is a representative of three experiments (n = 3 biologically independent experiments).

Subsequently, we selected a non-cysteine 9-mer antigen (NLVPMVATV), commonly known as pp65 viral antigen, derived from cytomegalovirus (CMV) and effectively presented by *HLA-A2*-encoded MHC-I^47^. We employed the SCT model to explore the reactivity of cysteines individually introduced at all positions of the pp65 antigen (pp65-C1-C9, **Fig. 5f**). The results revealed relatively low expressions of SCTs when cysteines were introduced at positions 2 and 9, likely due to the preference of *HLA-A2* encoded MHC-I for Leu, Met, or Ile at position 2 and Val, Leu, or Ile at position 9 (**Fig. 3c**). This suggests that introducing cysteines at these sites may disrupt proper complex folding, resulting in decreased protein expression. Notably, among cysteine-containing antigens showing similar protein expressions, MSD labeling patterns exhibited significant variations (**Fig. 5f**). pp65-C6 displayed the highest labeling despite lower protein expression compared to most other mutants. Even with low expression, pp65-C2 demonstrated strong probe labeling. In contrast, pp65-C3 showed minimal labeling despite having the highest expression level among these mutants. These findings indicate that cysteines on MHC-I-bound antigens may exhibit distinct reactivities, and the sulfonated maleimide probes are capable of effectively mapping these reactive cysteines using the SCT model.

### Reactivity-based antigen profiling for the discovery of reactive cysteines present on MHC-I-bound antigens

Cysteine-directed ABPP platforms have not only provided comprehensive maps of reactive cysteines across the human proteome, but also advanced investigations into electrophilic small molecule-cysteine interactions for novel ligand discovery^19, 48^. Building on this principle, we explored whether sulfonated maleimide probes could facilitate global mapping of reactive cysteines on MHC-I-bound antigens. To this end, we developed a chemical proteomics strategy wherein cells are treated with the MS-DTB probe, followed by cell lysis, streptavidin enrichment, and MS analysis to identify reactive cysteines present on MHC-I-bound antigens (**Fig. 6a**). To validate this platform, we initially overexpressed KRAS-G12C in BV173 parental and *HLA* knockout cells, then treated them with MS-DTB. Subsequent proteomic analysis revealed the identification of MS-DTB-modified KRAS-G12C neoantigen only in BV173 parental cells, but not in *HLA* knockout cells (**Fig. 6b**), indicating the feasibility of utilizing this approach to further profile reactive antigen cysteines in an untagged manner.

**Fig. 6.**
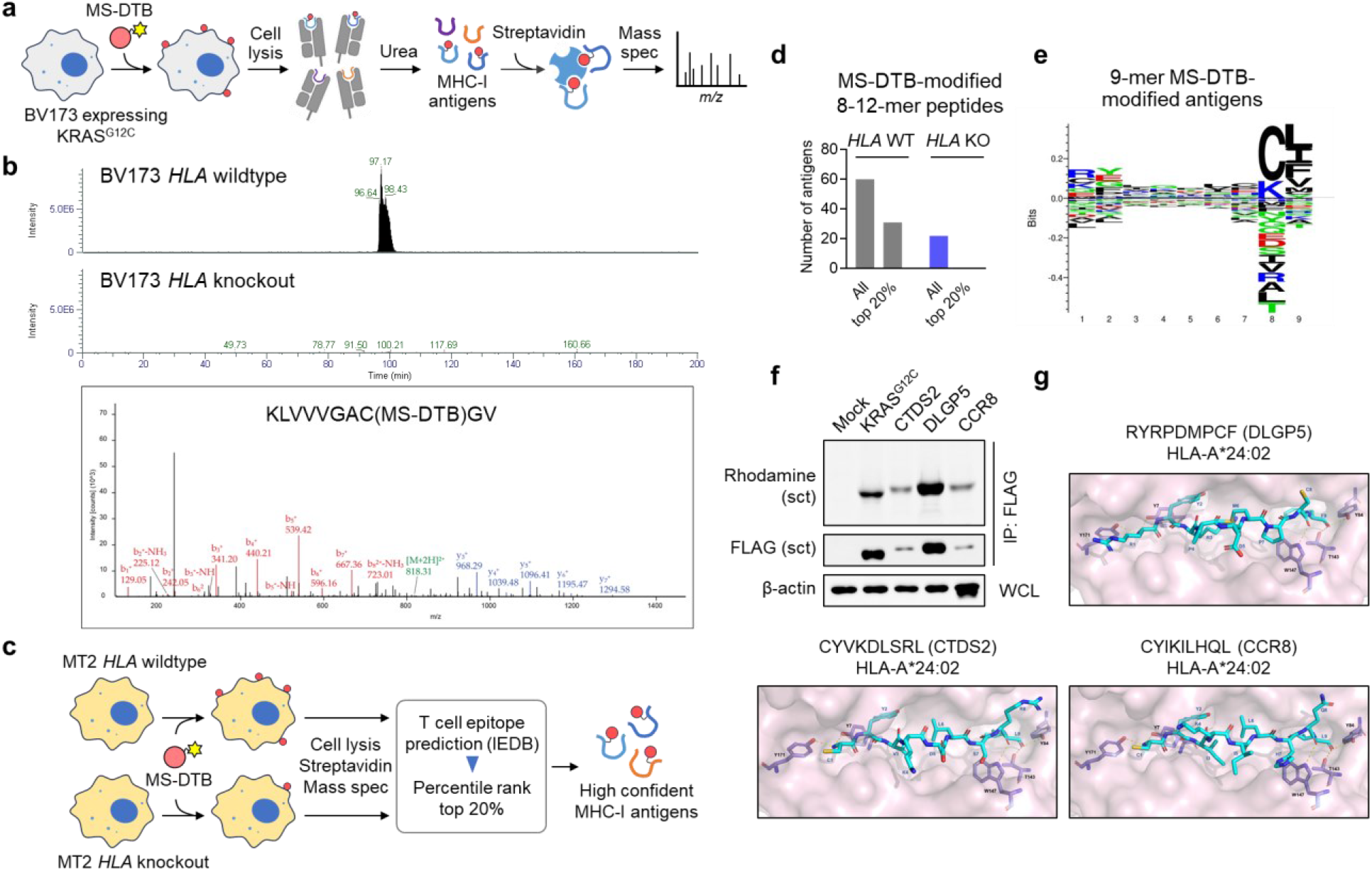
Reactivity-based antigen profiling for the discovery of reactive cysteines present on MHC-I-bound antigens. **a.** Schematic representation of employing MS-DTB to identify the KRAS-G12C neoantigen. **b**. MS-DTB-modified KRAS-G12C neoantigen was identified in BV173 wildtype, but not *HLA* knockout cells. The result is a representative of three experiments (n = 3 biologically independent experiments). **c**. Schematic representation of the reactivity-based antigen profiling workflow. **d**. Number of 8-12-mer MS-DTB-modified peptides identified in MT2 wildtype and *HLA* knockout cells. The result is a representative of three experiments (n = 3 biologically independent experiments). **e**. Motif analysis of 9-mer MS-DTB-modified peptides. **f**. Validation of three MS-DTB-modified peptides using SCTs. The result is a representative of two experiments (n = 2 biologically independent experiments). **g**. Modeling studies indicate that the cysteines in the three MS-DTB-modified peptides are solvent exposed in the context of forming pMHC-I complexes.

Subsequently, we implemented this reactivity-based antigen profiling strategy in MT2 parental and *HLA* knockout cells, aiming to identify new reactive cysteine-containing MHC-I antigens. One limitation of our initial platform is the inability of MS-DTB to differentiate MHC-I-bound antigens from peptides originating from other sources. To mitigate this limitation, we incorporated three filters into the workflow (**Fig. 6c**): 1) restriction to 8-12-mer peptides in the search engine to exclude longer peptides from the pMHC-II complex; 2) utilization of a T cell epitope prediction algorithm to retain only top-ranked peptides; and 3) selection of probe-modified peptides identified exclusively in MT2 parental cells, excluding peptides identified in *HLA* knockout cells. Using these filters, 60 MS-DTB-modified 8-12-mer peptides were identified in MT2 parental cells, with 31 falling within the top 20% ranking range using the T cell epitope prediction algorithm in IEDB (**Fig. 6d** and **Supplementary Table 6**). Motif analysis of 9-mer MS-DTB-modified peptides revealed a distribution pattern consistent with the HLA alleles (HLA-A*24:01 and HLA-B*40:01) in MT2 cells, suggesting that these probe-modified peptides likely originate from the pMHC-I complex (**Fig. 6e**). Conversely, in MT2 *HLA* knockout cells, all 22 MS-DTB-modified 8-12-mer peptides are below the top 20% ranking range, indicating that these peptides likely represent background noise. Notably, the preferred MS-DTB labeling sites at positions 1, 7, and 8 are consistent with findings from immunopeptidomics (**Fig. 3d,e**), further supporting the hyperreactivity of cysteines at these positions. Subsequently, we employed SCTs of HLA-A*24:02 to validate three MS-DTB-enriched antigens: CTDS2 (CYVKDLSRL), DLGP5 (RYRPDMPCF), and CCR8 (CYIKILHQL). In-gel fluorescence analysis demonstrated effective labeling of all three antigens by the MSD probe (**Fig. 6f**). Moreover, a modeling study using HLA-A*24:02 revealed that the anchor residues at positions 2 and 9 of all three antigen peptides fit into the B- and F-pockets, respectively (**Fig. 6g**). In the case of DLGP5, where cysteine resides at position 8, the phenylalanine at anchoring position 9 is buried within the F-pocket and forms hydrogen bonds with Y84 and T143 in MHC-I. The carbonyl group of C8 is fixed with the NH in W147 through a hydrogen bond, which causes the cysteine at position 8 to be solvent exposing. As for CTDS2 and CCR8 antigens, where cysteine is situated at position 1, the hydroxyl groups of Y7 and Y171 on MHC-I orient toward the A-pocket, forming hydrogen bonds with the N-terminal NH. Consequently, this arrangement results in the thiol of the cysteine residue being exposed to the solvent.

## Discussion

In this study, we introduce a platform that employs a cell-impermeable cysteine-reactive probe to map reactive cysteines on MHC-I-bound antigens. Our investigations revealed a previously overlooked phenomenon: not all cysteines on MHC-I-bound antigens are intrinsically reactive – they exhibit various degrees of reactivity. This variability may stem from their positions on antigens and the unique conformations adopted by individual pMHC-I complexes. These conformational differences could impact the pKa value of cysteines, resulting in varied cysteine reactivity. Harnessing cysteine reactivity is a fundamental strategy in covalent drug and chemical probe discovery^14^. Our findings suggest potential opportunities for targeting reactive cysteines on MHC-I antigens with electrophilic small molecules or biomolecules, thereby reshaping pMHC-I complexes on the cell surface for therapeutic interventions. More specifically, our findings, which highlight hyperreactive cysteines at position 8 on 9-mer antigens from two HLA alleles (HLA-A*24:02/HLA-B*40:01 and HLA-A*02:01/HLA-B*18:01) (**Fig. 3d,e**), suggest the potential of combining a moderate electrophilic warhead with tailored non-covalent moieties to recognize protein pockets within the TCR-pMHC-I complex, enabling selective antigen targeting. Furthermore, oxidative stress has been shown to promote the generation of post-translational neoantigens, influencing antigen-specific immunity dynamics^49^. For example, viral infection communicates with cytotoxic T cells through the *S*-glutathionylation occurring on the cysteines of MHC-I-presented viral peptides^50^. These neoantigens may arise from the proteasomal degradation of post-translationally modified proteins or the direct sensing of hyperreactive cysteines on surface-located antigens in the oxidative environment. Our platform has the potential to effectively map changes in cysteines within the MHC-I immunopeptidome, providing crucial insights into antitumor and antiviral immune responses.

Recent studies have demonstrated a unique mechanism for covalent drugs capable of hijacking antigen presentation pathways, resulting in the presentation of drug-modified antigens on MHC-I^16, 17^. These “chemical neoantigens” have the potential to open new avenues in cancer immunotherapy. As demonstrated by the authors, antibodies targeting these “chemical neoantigens” can be integrated into Bispecific T cell Engagers (BiTEs) to recruit cytotoxic T cells. A fundamental question following these studies is: what other covalent drugs or electrophilic small molecules have the potential to generate these chemical neoantigens? Our platform can facilitate the discovery of these molecules in various ways. First, the compatibility of sulfonated maleimide probes in ELISA assays enables the high-throughput measurement of various small molecules, biomolecules, and physiological conditions that could induce the formation of chemical neoantigens by hindering the engagement of sulfonated maleimide probes within pMHC-I complexes (**Fig. 2g, h**). Second, utilizing sulfonated maleimide probes in immunopeptidomics and reactivity-based antigen profiling provides a way to identify the sequences of these chemical neoantigens. Through multiplexed proteomics, the discovery of chemical neoantigens can be extended across multiple cell types and states.

Nonetheless, we recognize the limitations of our initial platform. One such limitation is that sulfonated maleimide probes do not inherently label MHC-I-bound antigens exclusively. They may also label MHC-II-bound longer antigens and a small fraction of extracellular proteins containing the reduced form of cysteines in specific physiological contexts. Therefore, at the current stage, incorporating a pairwise cellular model with *HLA* knockout and integrating a T cell epitope prediction algorithm to retain only the top-ranked peptides would be beneficial in narrowing down the observed effect to MHC-I-associated antigens.

## Acknowledgement.

We gratefully acknowledge the support of the Ono Pharma Foundation. We thank the Robert H. Lurie Comprehensive Cancer Center of Northwestern University for the use of the Flow Cytometry Core Facility.

---

**Fig. S1.**
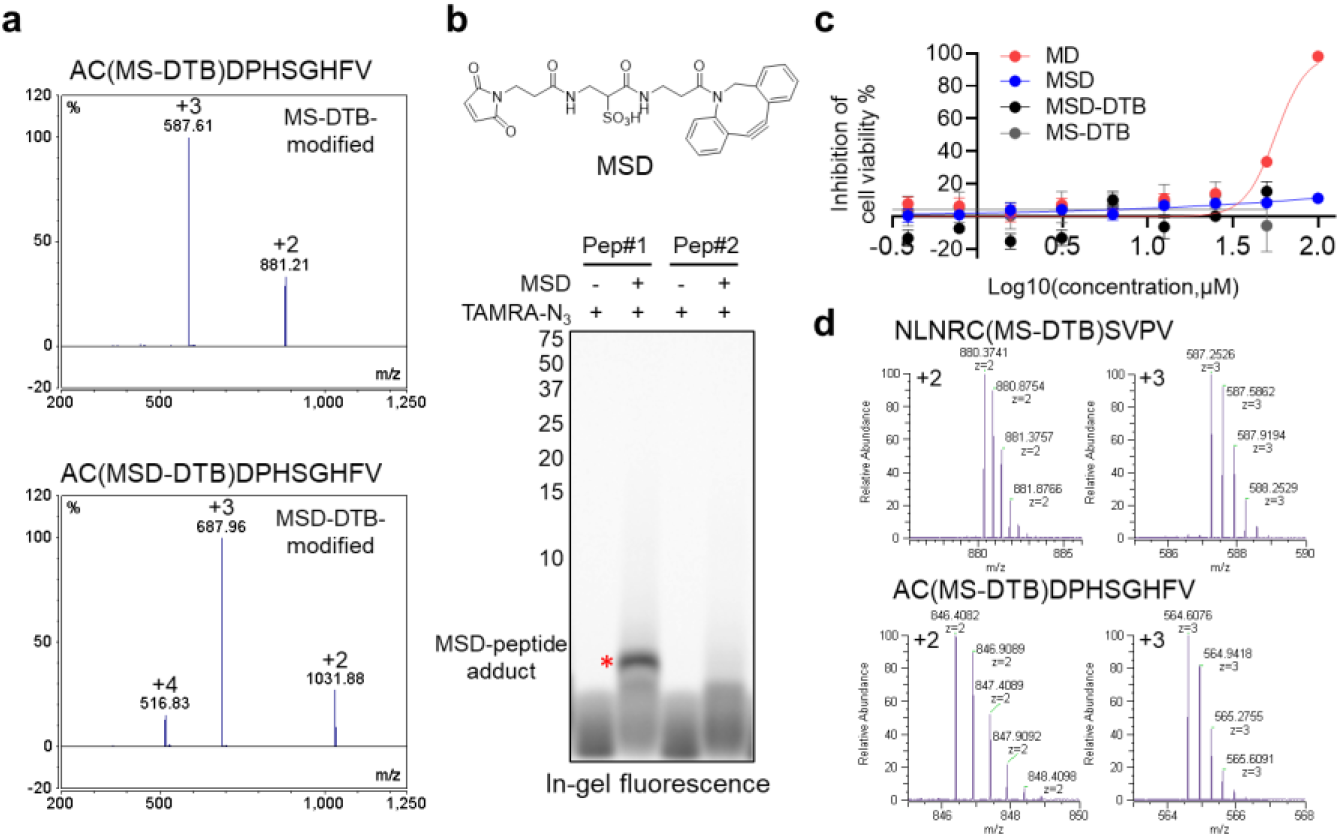
Development of chemical probes for mapping cysteines within the immunopeptidome. **a**. MS1 spectra of MSD-DTB-peptide and MS-DTB-peptide adducts from the LC-MS analysis in Fig. 1c. **b**. In-gel fluorescence analysis revealed that MSD probe selectively labels cysteine-containing peptide. Peptides were analyzed through tricine gels. The result is a representative of two experiments (n = 2 biologically independent experiments). **c**. Inhibition of HEK293T cell viability by MD, MSD, MSD-DTB and MS-DTB probes (72 hours). Data represents mean values ± SEM (n = 3 biologically independent experiments). **d**. MS1 spectra of two peptides modified by MS-DTB in the orbitrap mass analyzer. Both modified peptides were doubly and triply charged and showed normal isotopic envelope profiles.

**Fig. S2.**
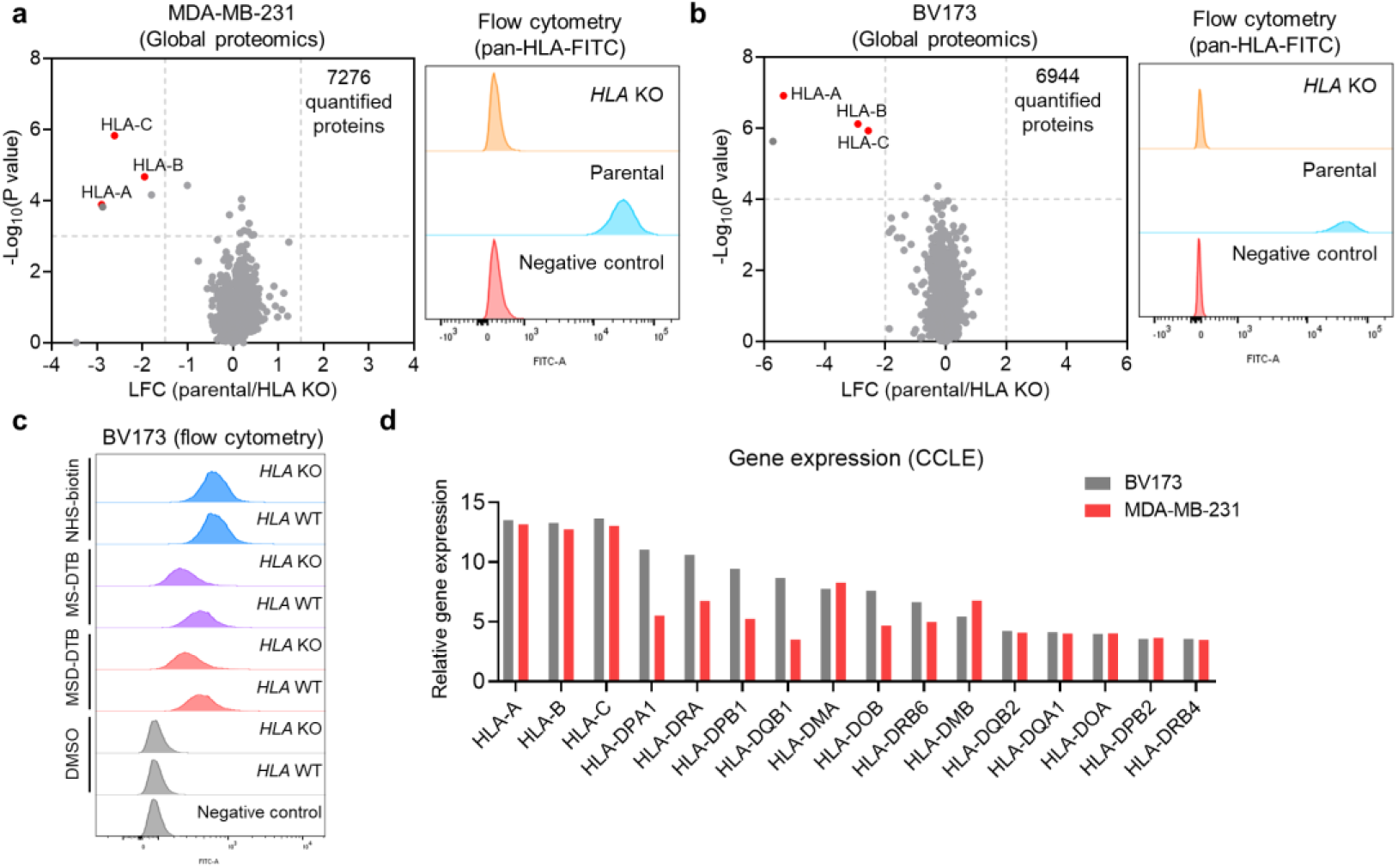
Sulfonated maleimide probes modify cysteines present on MHC-I-bound antigens. **a**. Quantitative global proteomics and flow cytometry studies confirmed the knockout of *HLA-A*, *HLA-B*, and *HLA-C* in MDA-MB-231 cells (n = 3 biological independent samples for global proteomics, n = 3 biological independent samples for flow cytometry). **b**. Quantitative global proteomics and flow cytometry studies confirmed the knockout of *HLA-A*, *HLA-B*, and *HLA-C* in BV173 cells (n = 3 biological independent samples for global proteomics, n = 3 biological independent samples for flow cytometry). **c**. Flow cytometry histograms comparing parental and *HLA* knockout cells treated with MSD-DTB, MS-DTB, and sulfonated NHS-biotin probes. **d**. mRNA expression levels of genes encoding for MHC-I and MHC-II in BV173 and MDA-MB-231 cells. Data was obtained from Cancer Cell Line Encyclopedia (CCLE) (https://sites.broadinstitute.org/ccle/).

**Fig. S3.**
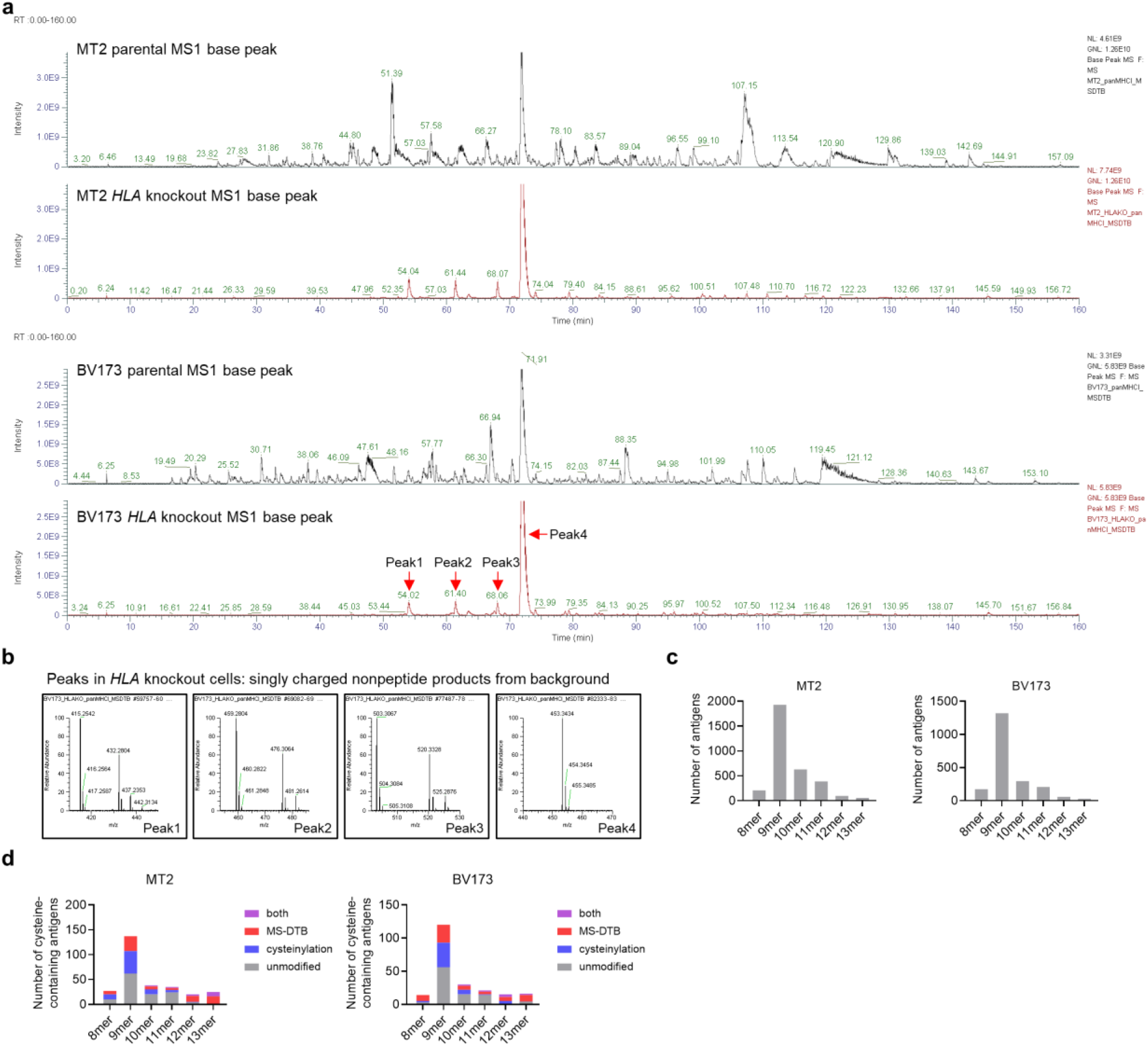
MHC-I immunopeptidomics studied in MT2 and BV173 cells treated with MS-DTB. **a.** MS1 base peak chromatograms of immunopeptidomics comparing wildtype versus *HLA* knockout in MT2 and BV173 cells. The result is a representative of two experiments (n = 2 biologically independent experiments). **b**. MS1 spectra of four representative peaks identified in BV173 *HLA* knockout cells. **c**. The number of 8-13-mer antigens in MT2 and BV173 cells. **d**. The number of 8-13-mer cysteine-containing antigens in MT2 and BV173 cells.

**Fig. S4.**
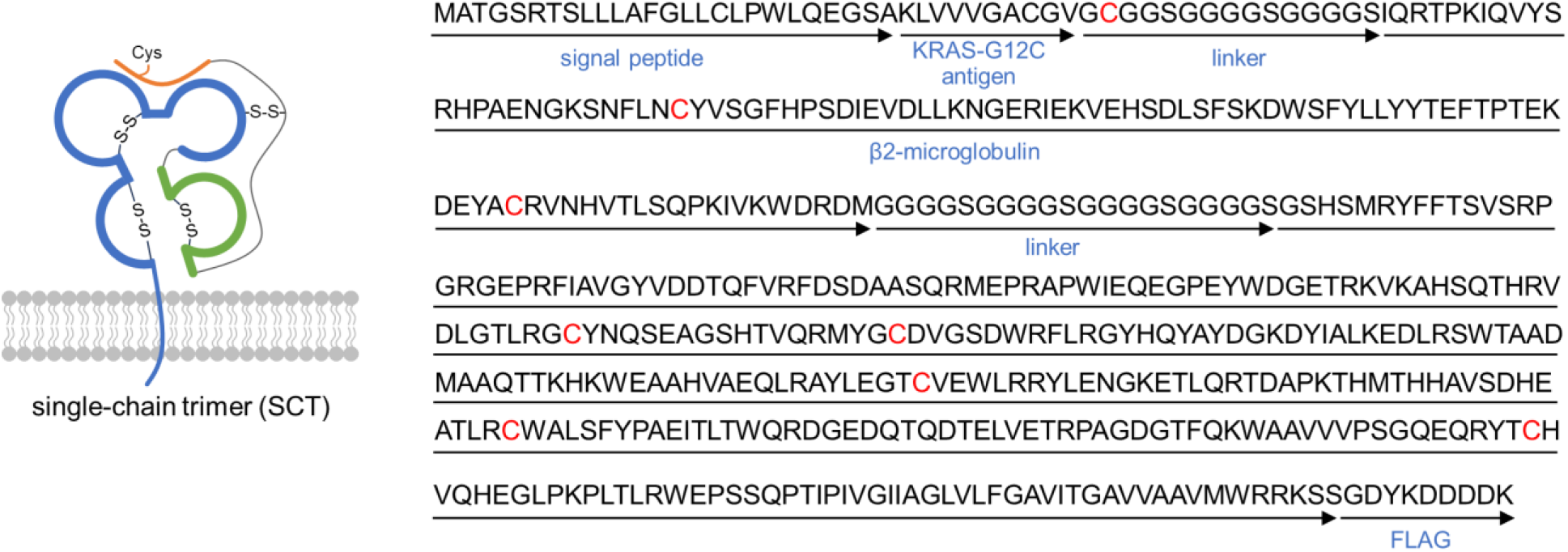
The translated single-chain peptide sequence from the SCT construct of *HLA-A2*/KRAS-G12C. Cysteines are highlighted in red.

## Materials and Methods

### Reagents

The anti-FLAG HRP antibody (clone M2, cat#: A8592) and anti-FLAG affinity gel (clone M2, cat#: A2220) were purchased from Sigma-Aldrich. The anti-β-Actin antibody (clone#: C4, cat#: sc-47778) was purchased from Santa Cruz Biotechnology. Streptavidin-HRP (cat#: 3999), anti-GAPDH (clone 14C10, cat#: 3683), anti-MHC Class I (clone EMR8-5, cat#: 88274) and anti-PSMB9 (clone E7J1L, cat#: 87667) were purchased from Cell Signaling Technology. Puromycin (cat#: ant-pr-1) was purchased from InvivoGen. InVivoMAb anti-human MHC Class I (HLA-A, HLA-B, HLA-C) (clone W6/32, cat#: BE0079) for ELISA assay was purchased from Bio X Cell. Ultra-LEAF anti-human HLA-A,B,C antibody (clone W6/32, cat#: 311448) for immunopeptidomics was purchased from BioLegend. Polyethylenimine (PEI, MW 40,000, cat#: 24765-1) was purchased from Polysciences, Inc. Tetramethylrhodamine (TAMRA) azide (cat#: T10182), enzyme-linked chemiluminescence (ECL) (cat#: 32106) western blotting detection reagents, QuantaBlu fluorogenic peroxidase substrate kit (cat#: 15169), Streptavidin agarose (cat#: 20349), Streptavidin-FITC (cat#: 11-4317-87), HLA-ABC FITC antibody (clone W6/32, cat# MA5-44095), beta-2-microglobulin HRP (clone B2M-01, cat#: MA1-19679), and Tandem Mass Tag (TMT) isobaric label reagent (cat#: 90066 for TMTsixplex and cat#90406 for TMT10plex) were purchased from Thermo Scientific. FuGene 6 (cat#: E2692) transfection reagent and sequencing grade modified trypsin (cat#: V5111) were purchased from Promega. Cas9 endonuclease was purchased from Integrated DNA Technologies. pMHC-I HLA-A2-KRASG12C was purchased from ProImmune. Maleimide-sulfonate-dibenzocyclooctyne (MSD), Maleimide-dibenzocyclooctyne (MD), Maleimide-sulfonate-PEG4-dibenzocyclooctyne (MSD4), and N-hydroxysuccinimide (NHS)-sulfonate-biotin were purchased from BroadPharm.

### Cell lines

HEK293T and MDA-MB-231 cells were obtained from ATCC. BV173 cells were obtained from CLS Cell Lines Service. MT2 cells were obtained from Thermo Scientific. HEK293T and MDA-MB-231 cells were cultured in Dulbecco’s Modified Eagle Medium (DMEM, Corning) with 10% (v/v) fetal bovine serum (FBS, Omega Scientific) and L-glutamine (2mM, Gibco). BV173 and MT2 cells were cultured in RPMI 1640 (Corning) with 10% (v/v) fetal bovine serum (FBS, Omega Scientific) and L-glutamine (2mM, Gibco). All the cell lines were tested negative for mycoplasma contamination.

### Generation of CRISPR-Cas9-mediated HLA knockout cells

BV173, MT2 and MDA-MB-231 cells with HLA-A,B,C CRISPR-Cas9 knockout were generated through electroporation of Cas9-sgRNA ribonucleoprotein (RNP) complex using 4D-Nucleofector (Lonza Bioscience). Three sgRNAs targeting HLA gene (HLA sgRNA#1: CGGCTACTACAACCAGAGCG; HLA sgRNA#2: AGATCACACTGACCTGGCAG; HLA sgRNA#3: AGGTCAGTGTGATCTCCGCA) were mixed for the electroporation.

### Cloning and mutagenesis

Human KRAS4A-G12C cDNAs with N-terminal FLAG tag and all single-chain trimers were purchased as gene block from Integrated DNA Technologies and cloned into pCDH-CMV-MCS-EF1-Puro vector via NehI and BamHI sites. pp65 mutants were generated using Q5 site-directed mutagenesis kit (New England Biolabs).

### Generation of KRAS4A-G12C stably expressed cells by lentivirus transduction

Lentivirus containing FLAG-KRAS4A-G12C were generated by co-transfection of FLAG-KRAS4A-G12C, psPAX2 and pMD2.G into HEK293T cells using FuGene 6 transfection reagent. Medium containing lentiviral particles were collected 48 hours post transfection, filtered with 0.45 µM Millex-HV sterile syringe filter unit (MilliporeSigma), and used to transduce BV173 cells in the presence of 10 µg/mL polybrene. 48 hours post transduction, puromycin (2 µg/mL) was added and incubated with the cells for 7 days.

### Cell lysis and Western blot

Cells were lysed utilizing RIPA lysis buffer (Thermo Scientific) comprising 25 mM Tris-HCl, pH 7.6, 150 mM NaCl, 1% Nonidet P40 (NP-40), 1% sodium deoxycholate, and 0.1% sodium dodecyl sulfate (SDS). Before usage, the lysis buffer was supplemented with the cOmplete protease inhibitor cocktail (Roche). The cell suspension underwent sonication through 5 cycles at 40% power for 4 pulses each. Subsequent to sonication, the resultant mixture underwent centrifugation at 16,000g for 10 minutes at 4°C to acquire the supernatant. The protein concentration in the supernatant was determined employing the DC assay (Bio-Rad). The protein lysate was combined with Laemmli sample buffer (Bio-Rad) and heated at 95°C for 5 minutes. Proteins were analyzed using 4-20% Novex Tris-Glycine mini gels (Invitrogen), followed by transfer onto a 0.2 µM polyvinylidene fluoride (PVDF) membrane (Bio-Rad). The PVDF membrane was incubated with 5% non-fat milk in TBST buffer (0.1% Tween 20, 20 mM Tris-HCl at pH 7.6, and 150 mM NaCl) for 1 hour at room temperature. Primary antibodies were diluted in 5% non-fat milk in TBST buffer and incubated with the membrane. Incubation durations were 1 hour at room temperature for FLAG and β-actin, and overnight at 4°C for others. Following primary antibody incubation, the membrane underwent three washes with TBST buffer and was then incubated with a secondary antibody (diluted 1:5000 in 5% non-fat milk in TBST) for 1 hour at room temperature. After three additional washes with TBST buffer, the chemiluminescence signal on the membrane was developed using ECL western blotting detection reagent, and the resultant signal was captured using ChemiDoc MP (Bio-Rad).

### Immunoprecipitations

Cells were lysed in NP-40 lysis buffer (25 mM Tris-HCl, pH 7.4, 150 mM NaCl, 10% glycerol, 1% NP-40) supplemented with cOmplete protease inhibitor cocktail. The cell suspension was incubated on ice for 10 minutes. Following this, the mixture was centrifuged at 16,000 g for 10 minutes at 4°C, and the resulting supernatant was collected for use in immunoprecipitation. For immunoprecipitation, FLAG affinity gel (25 µL slurry per sample) was added to the protein lysates and rotated at 4°C for 2 hours. The affinity gel was then washed four times with immunoprecipitation washing buffer comprising 0.2% NP-40, 25 mM Tris-HCl at pH 7.4, and 150 mM NaCl. Subsequently, the affinity gel was mixed with Laemmli sample buffer and heated at 95°C for 10 minutes. The resulting supernatant, containing the eluted proteins, was collected and utilized for subsequent western blot analysis.

### Global proteomics analysis

Cells were lysed in 100 µL of PBS using sonication (10 pulses at 40% intensity, 3 rounds). Protein concentration was determined via a DC assay. Next, 100 µg of proteins in 100 µL of lysis buffer were denatured with 8 M urea. For reduction, 5 µL of 200 mM DTT stock solution in water was added, and the mixture was heated to 65°C for 15 minutes. Alkylation was achieved by adding 5 µL of 400 mM iodoacetamide stock solution in water and incubating in the dark at 37°C for 30 minutes. Proteins were then precipitated by adding 600 µL of MeOH, 200 µL of CHCl_3_, and 500 µL of water. After precipitation, protein pellets were washed with 1 mL of MeOH. The resulting protein pellets were solubilized in 160 µL of EPPS buffer (200 mM). Subsequently, 2 µg of LysC was added to each sample, and digestion was carried out at 37°C for 2 hours. This was followed by the addition of 5 µg of trypsin to each sample for another round of digestion, allowed to proceed at 37°C for 12 hours. For TMT labeling, 12.5 µg of resulting peptides in 35 µL of EPPS buffer were utilized. To each sample, 9 µL of CH_3_CN was added, followed by TMT tags (3 µL per sample). The samples were then incubated at room temperature for 1 hour. The TMT labeling reaction was quenched by adding 6 µL of a 5% hydroxylamine solution, followed by the addition of 2.5 µL of formic acid. The samples were pooled and separated into 12 distinct fractions using the Thermo Vanquish UHPLC fractionator. These fractions were analyzed on an Orbitrap Eclipse Tribrid mass spectrometer coupled with a Vanquish Neo UHPLC system. Peptides were injected onto an EASY-Spray HPLC column (C18, 2 µm particle size, 75 µm inner diameter, 250 mm length) and eluted at a flow rate of 0.25 µL/min, following a gradient: 5% buffer B (80% acetonitrile with 0.1% formic acid) in buffer A (water with 0.1% formic acid) from 0 to 15 minutes, 5% to 45% buffer B from 15 to 155 minutes, and 45% to 100% buffer B from 155 to 180 minutes. The parameters for the MS1 scan are: resolution 120,000, m/z range 375-1600, RF lens 30%, standard AGC target and auto maximum injection time. In the MS2 analysis, precursor ions were quadrupole-isolated (isolation window 0.7) and then subjected to HCD collision in the ion trap (standard AGC, collision energy 30%, maximum injection time 35 ms). Following each MS2 spectrum, synchronous precursor selection (SPS) enabled the selection of 10 MS2 fragment ions for MS3 analysis. These MS3 precursors were fragmented by HCD and analyzed using the Orbitrap (collision energy 55%, AGC 250%, maximum injection time 200 ms, resolution 60,000). The RAW data was analyzed using Proteome Discoverer 2.5.

### Cysteine-directed ABPP

Cells were lysed in PBS via sonication (10 pulses at 40% intensity, 3 rounds). The protein concentration was determined using a DC assay and adjusted to 1 mg/mL. Next, 500 µL of lysates were labeled with 100 µM IA-DTB or DBIA at room temperature for 1 hour. Protein precipitation was achieved by adding 500 µL of methanol and 100 µL of chloroform, followed by a methanol wash (1 mL). The resulting protein pellets were denatured using 90 µL of 9 M urea and 10 mM DTT in 50 mM tetramethylammonium bicarbonate. Alkylation was carried out using 50 mM iodoacetamide at 37°C for 30 minutes. Subsequently, 350 µL of 50 mM tetramethylammonium bicarbonate was added to each sample, followed by the addition of 2 µg of trypsin. Digestion was allowed to proceed at 37°C for 12 hours. Next, 50 µL of streptavidin-agarose beads were added to each sample, and the mixture was rotated at room temperature for 2 hours. The beads were washed three times with 1 mL of washing buffer consisting of 0.2% NP-40, 25 mM Tris-HCl pH 7.4, and 150 mM NaCl, followed by three washes with 1 mL of PBS, and two washes with 1 mL of water. Peptides were eluted using 300 µL of 50% acetonitrile containing 0.1% formic acid. The eluted peptides were subsequently dried using a SpeedVac vacuum concentrator. The subsequent steps of TMT labeling and LC-MS analysis were carried out following the methodology described in Global proteomics analysis.

### Immunopeptidomics

2 × 10^8^ BV173 or MT2 cells were lysed using 3 mL of lysis buffer (0.5% NP-40, 50 mM Tris pH 8.0, 150 mM NaCl, 1 mM EDTA, and protease inhibitor cocktail) by rotating at 4°C for 30 minutes. Following centrifugation at 18,000 g for 10 minutes, the supernatant was collected for enrichment using an anti-MHC antibody (W6/32, BioLegend) conjugated to Affi-gel 10 matrix (Bio-Rad, 2 mg of antibody per 100 µL of slurry per sample). Enrichment occurred over 4 hours of rotation at 4°C, followed by transfer to a Bio-spin column (Bio-Rad) for washing with 3 x 1 mL of lysis buffer, wash buffer 1 (50 mM Tris pH 8, 150 mM NaCl), wash buffer 2 (50 mM Tris pH 8, 400 mM NaCl), and wash buffer 3 (50 mM Tris pH 8). MHC-conjugated peptides were subsequently eluted using 1 mL of 1% trifluoroacetic acid in water. Peptide samples were desalted using a Sep-Pak C18 cartridge (Waters), dried via speedavac, and analyzed using an Orbitrap Eclipse Tribrid mass spectrometer coupled with a Vanquish Neo UHPLC system.

### Reactivity-based antigen profiling

10^8^ cells were treated with 50 µM of MS-DTB for 30 minutes. After treatment, the cells were washed twice with PBS and then harvested. Subsequently, the cells were lysed in 5 mL of lysis buffer containing 2M urea and 0.2% NP-40 in PBS using sonication (10 pulses at 40% intensity, 3 rounds). Following centrifugation at 18,000 g for 10 minutes, the supernatant was collected for enrichment using streptavidin agarose beads. The mixture was rotated at room temperature for 2 hours. The beads were then washed three times with 1 mL of washing buffer (0.2% NP-40, 25 mM Tris-HCl pH 7.4, and 150 mM NaCl), followed by three washes with 1 mL of wash buffer 1 (50 mM Tris pH 8, 150 mM NaCl), wash buffer 2 (50 mM Tris pH 8, 400 mM NaCl), PBS, and water. Peptides were eluted using 300 µL of 50% acetonitrile containing 0.1% formic acid. The eluted peptides were subsequently dried using a SpeedVac vacuum concentrator and desalted using a Sep-Pak C18 cartridge. The peptides were analyzed using an Orbitrap Eclipse Tribrid mass spectrometer coupled with a Vanquish Neo UHPLC system.

### Modeling study

The crystal structures of HLA-A*02:01 (2X4R) and HLA-A*24:02 (2BCK) from Protein Data Bank (X-ray structures with a resolution finer than 3.5 Å), KRAS^G12C^ neoantigen (KLVVVGACGV) and three MS-DTB-enriched antigens: CTDS2 (CYVKDLSRL), DLGP5 (RYRPDMPCF), and CCR8 (CYIKILHQL) were used for the modeling study. MHC-Fine, a refined AlphaFold model, was used for MHC-peptide complex prediction^51^. The final PDB files of the MHC-peptide complex were generated by running the inference code and datasets (https://bitbucket.org/abc-group/mhc-fine/src/main/). The figures were generated by PyMOL software. For each MHC protein, only the α1 and α2 domains were used.

The FASTA sequence of HLA-A*02:01: GSHSMRYFFTSVSRPGRGEPRFIAVGYVDDTQFVRFDSDAASQRMEPRAPWIEQEGPEYWDGETRKVKAHSQTHRVDLGTLRGYYNQSEAGSHTVQRMYGCDVGSDWRFLRGYHQYAYDGKDYIALKEDLRSWTAADMAAQTTKHKWEAAHVAEQLRAYLEGTCVEWLRRYLENGKETLQRT

The FASTA sequence of HLA-A*24:02: GSHSMRYFSTSVSRPGRGEPRFIAVGYVDDTQFVRFDSDAASQRMEPRAPWIEQEGPEYWDEETGKVKAHSQTDRENLRIALRYYNQSEAGSHTLQMMFGCDVGSDGRFLRGYHQYAYDGKDYIALKEDLRSWTAADMAAQITKRKWEAAHVAEQQRAYLEGTCVDGLRRYLENGKETLQRT

### Cell surface probe labeling by flow cytometry

Cells were seeded in non-treated 6-well plates and treated with 50 µM of reactivity probes for 30 minutes. Following treatment, cells were rinsed with PBS and suspended in flow cytometry buffer (1 mM EDTA, 25 mM HEPES pH 7.0, 1% FBS in PBS) within Eppendorf tubes. Subsequently, Streptavidin-FITC or HLA-ABC-FITC antibody (diluted 1:50) was added, and the cells were rotated at room temperature for 30 minutes. Afterward, the cells were washed with PBS and resuspended in flow cytometry buffer. FITC fluorescence on the cell surface was quantified using a BD LSRFortessa Cell Analyzer, and the resulting data were analyzed utilizing FlowJo software.

### LC-MS analysis of probe-peptide adduct

Peptides were synthesized by GenScript. 100 µM peptide and 100 µM compound were incubated in water for 30 minutes and analyzed by Thermo Vanquish UHPLC coupled to ISQ EC Single Quadrupole Mass Spectrometer. Peptide and probe-peptide adduct were separated on Gemini C18 column (Phenomenex, 5 µm, 50 x 4.6 mm) at a flow rate of 1 mL/min, following the gradient: 0 to 95% buffer C (acetonitrile with 0.1% formic acid) in buffer A (water with 0.1% formic acid) from 0 to 13 minutes, and 95% buffer C in buffer A from 13 to 20 minutes. 210 nm wavelength was used to monitor the peaks. A full scan from m/z 200-1250 with positive mode was used to analyze unmodified peptide and probe-peptide adduct.

### ELISA assay

Nunc MaxiSorp 384-well plates (black) were coated overnight with 50 µL of the anti-heavy chain antibody W6/32 at a concentration of 5 µg/mL in PBS. Following coating, the plates were washed twice with PBS (100 µL) and blocked with 3% BSA in PBS (120 µL) at room temperature for 1 hour. Subsequently, the plates were washed three times with 0.05% Tween-20 in PBS (PBST) (100 µL each wash). BV173, MT2 parental, and *HLA* knockout cells were treated with 50 µM of MS-DTB for 30 minutes, then harvested and lysed in NP-40 lysis buffer with protease inhibitor cocktail. The protein concentration was adjusted to 1 mg/mL, and 50 µL of total lysates were added to each well. Plates were incubated at 4°C for 4 hours, followed by three washes with 1% BSA in PBS (100 µL each wash). Next, 50 µL of either 1 µg/mL anti-beta-2-microglobulin HRP conjugate solution or Streptavidin-HRP (diluted 1:1000) in 1% BSA PBS was added to each well. Plates were incubated with shaking at room temperature for 1 hour. The plates were washed three times with PBST and three times with PBS (100 µL each wash). 50 µL of the HRP substrate QuantaBlu was added, and fluorescence was measured using the CLARIOstar Plus microplate reader (BMG Labtech).

### In-gel fluorescence

HEK293T cells were transfected with SCTs using PEI transfection reagent. Following a 24-hour incubation, the cells were treated with 50 µM of MSD for 30 minutes. Subsequently, cells were harvested by centrifugation at 500 g for 5 minutes and then lysed in NP-40 lysis buffer containing protease inhibitor cocktail. The resulting lysate was subjected to immunoprecipitation with anti-Flag affinity gel at 4°C for 2 hours. The affinity gel was washed three times with immunoprecipitation washing buffer and re-suspended in 18 µL of PBS. For the click chemistry reaction, the following reagents were added: 0.8 µL of 1.5 mM TAMRA azide solution in DMSO, 1.2 µL of 10 mM TBTA solution in 4:1 tBuOH:DMSO, 1 µL of 40 mM CuSO_4_ solution in H_2_O, and 1 µL of 40 mM TCEP solution in H_2_O. The reaction proceeded at room temperature for 1 hour. Subsequently, Laemmli sample buffer was added and heated at 95 °C for 10 minutes. Following centrifugation at 15,000 g for 2 minutes, the supernatant was collected, and the samples were resolved by 4-20% Novex Tris-Glycine mini gels. In-gel fluorescence signals were recorded using the ChemiDoc MP system.

### Cell viability assay

Cells were plated in a 96-well clear bottom white plate (Corning) at a density of 5,000 cells per well in 100 µL of DMEM medium and incubated for 24 hours. Subsequently, the cells were treated with varying concentrations of compounds in 100 µL of DMEM medium for additional 72 hours. Following treatment, 50 µL of Cell Titer Glo reagent (Promega) was added to each well and incubated for 10 minutes at room temperature. Luminescence was measured using CLARIOstar Plus microplate reader (BMG Labtech).

### Statistical analysis

Quantitative data were depicted using scatter plots, displaying the mean accompanied by the standard error of the mean (SEM) represented as error bars. Differences between two groups were assessed using an unpaired two-tailed Student’s t-test. Significance levels were denoted as follows: \**P* < 0.05, \*\**P* < 0.01, and \*\*\**P* < 0.001. Statistical significance was defined for *P* values < 0.05.

### Synthetic procedures

Synthesis of maleimide-sulfonate-DTB (MS-DTB)

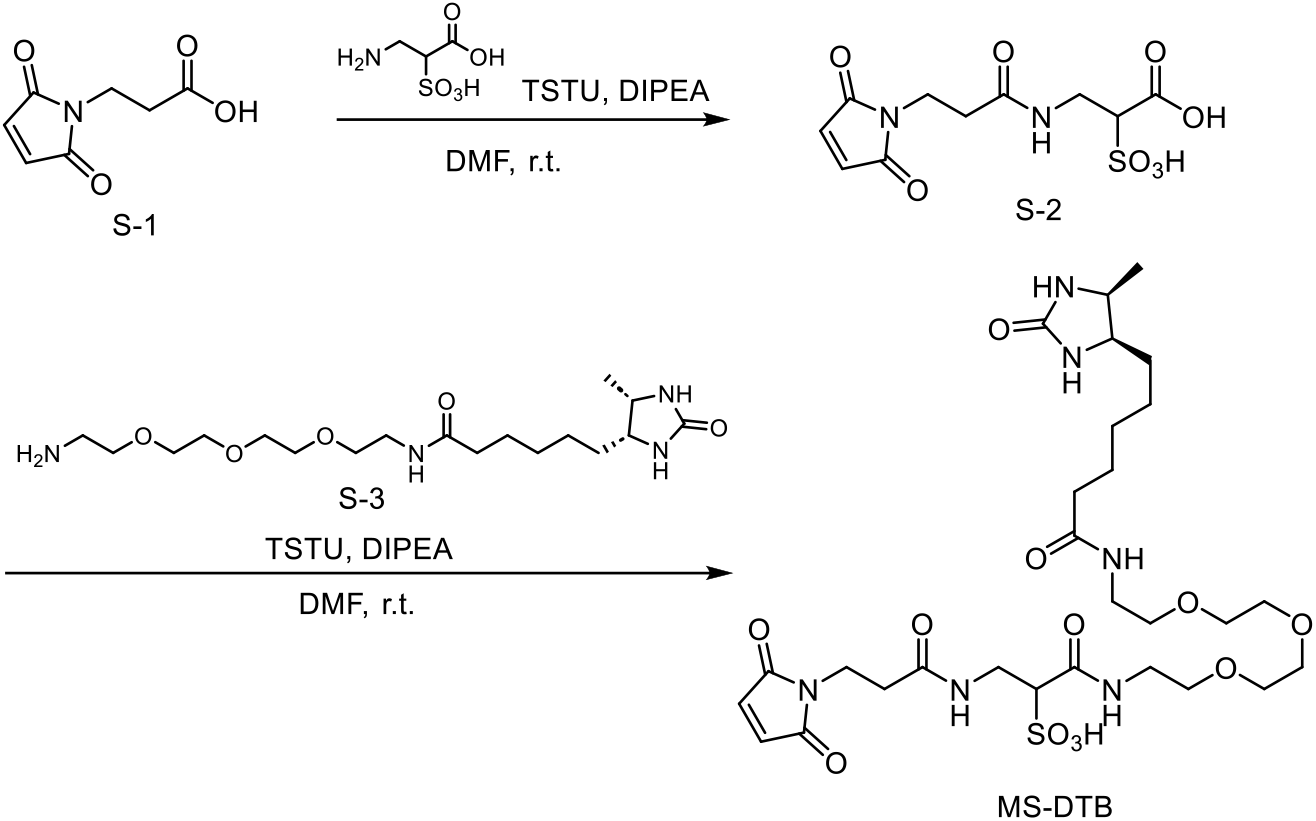

### Step 1

3-Maleimidopropionic acid S-1 (50 mg, 0.30 mmol, 1.0 eq) and TSTU (117 mg, 0.39 mmol, 1.3 eq) was dissolved in DMF (1 mL). Then DIPEA (149 μL, 0.90 mmol, 3.0 eq) was added and stirred for 1 h at room temperature. After the formation of succinate, 3-amino-2-sulfopropanoic acid (75 mg, 0.45 mmol, 1.5 eq) was added and stirred overnight. Upon completion, the reaction was concentrated, and the resulting residue was purified by flash chromatography to provide S-2 as a colorless oil (61 mg, 0.19 mmol, 63%).

**^1^H NMR** (400 MHz, CD_3_OD) δ 6.79 (s, 2H), 3.79 – 3.70 (m, 5H), 2.46 (t, *J* = 6.8 Hz, 2H).

### Step 2

S-3 was prepared as reported previously^52^. S-2 (10 mg, 0.031 mmol, 1.0 eq) and TSTU (12 mg, 0.040 mmol, 1.3 eq) was dissolved in DMF (1 mL). Then DIPEA (15 μL, 0.093 mmol, 3.0 eq) was added and stirred at room temperature for 1 hour. After the formation of succinate, S-3 (18 mg, 0.047 mmol, 1.5 eq) was added and stirred overnight. Upon completion, the reaction was concentrated, and the resulting residue was purified by flash chromatography to provide MS-DTB as a colorless oil (11.2 mg, 0.016 mmol, 52%).

**^1^H NMR** (500 MHz, CD_3_OD) δ 6.80 (s, 2H), 3.85 – 3.78 (m, 3H), 3.77 – 3.68 (m, 4H), 3.67 – 3.60 (m, 9H), 3.58 (t, *J* = 6.0 Hz, 2H), 3.55 (t, *J* = 5.5 Hz, 3H), 3.47 – 3.39 (m, 2H), 3.38 – 3.35 (m, 2H), 2.45 (t, *J* = 7.0 Hz, 3H), 2.22 (t, *J* = 7.5 Hz, 3H), 1.63 (p, *J* = 7.5 Hz, 3H), 1.53 – 1.48 (m, 2H), 1.47 – 1.35 (m, 3H), 1.11 (d, *J* = 6.0 Hz, 3H).

**^13^C NMR** (126 MHz, CD_3_OD) δ 176.37, 172.85, 172.22, 169.06, 166.23, 135.51, 79.51, 71.61, 71.39, 71.31, 70.62, 70.39, 66.09, 57.43, 52.73, 40.66, 40.40, 39.64, 36.94, 35.78, 35.40, 30.76, 30.24, 27.19, 26.87, 15.68.

**HRMS** (ESI+) m/z calcd for C_28_H_47_N_6_O_12_S^+^ [M+H]^+^: 691.2967, found 691.2956.

Synthesis of maleimide-sulfonate-dibenzocyclooctyne-DTB (MSD-DTB)

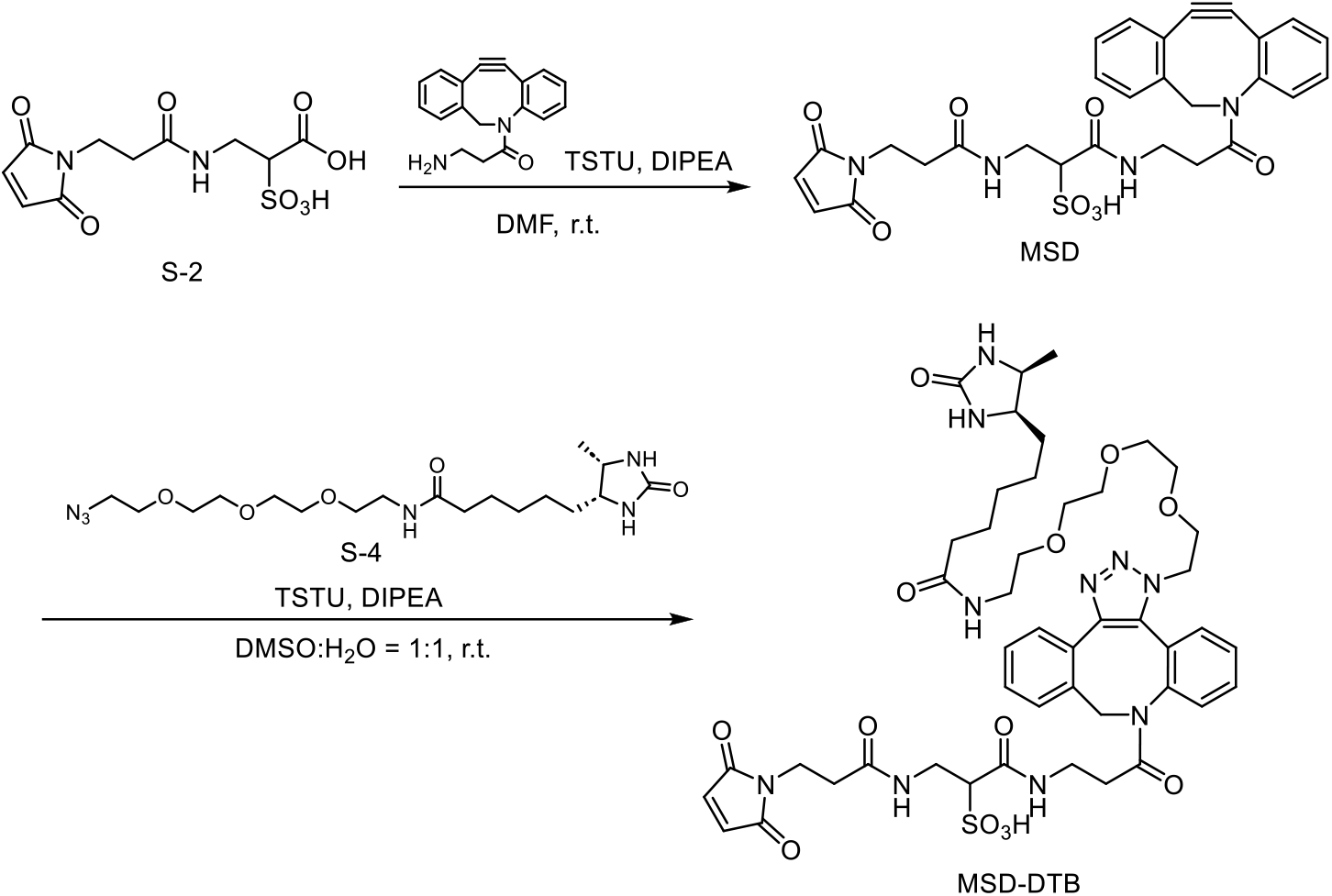

### Step 1

S-2 (40 mg, 0.125 mmol, 1.0 eq) and TSTU (49 mg, 0.163 mmol, 1.3 eq) was dissolved in DMF (1 mL). Then DIPEA (149 μL, 0.90 mmol, 3.0 eq) was added and stirred for 1 h at room temperature. After the formation of succinate, dibenzocyclooctyne-amine (52 mg, 0.188 mmol, 1.5 eq) was added and stirred overnight. Upon completion, the reaction was concentrated, and the resulting residue was purified by flash chromatography to provide MSD as a white powder (67 mg, 0.116 mmol, 71%).

**^1^H NMR** (500 MHz, CD_3_OD) δ 7.82 – 7.24 (m, 8H), 6.79 – 6.78 (m, 2H), 5.15 (dd, *J* = 14.0, 4.5 Hz, 1H), 3.77 – 3.56 (m, 6H), 3.43 – 3.37 (m, 1H), 3.28 – 3.24 (m, 1H), 2.65 – 2.57 (m, 1H), 2.44 – 2.36 (m, 2H), 2.07 – 1.98 (m, 1H).

### Step 2

S-4 was prepared as reported previously^53^. MSD (30 mg, 0.052 mmol, 1.0 eq) and S-4 (22 mg, 0.052 mmol, 1.0 eq) was dissolved in DMSO:H_2_O (1:1, 1 mL) and stirred at room temperature for 1 hour. Upon completion, the reaction was concentrated, and the resulting residue was purified by flash chromatography to provide MSD-DTB as a white powder (29.6 mg, 0.030 mmol, 58%).

**^1^H NMR** (500 MHz, CD_3_OD) δ 7.72 – 7.25 (m, 8H), 6.82 – 6.80 (m, 2H), 6.02 – 5.96 (m, 1H), 4.71 – 4.46 (m, 3H), 4.19 – 4.17 (m, 1H), 3.93 – 3.89 (m, 1H), 3.82 – 3.42 (m, 1H), 3.34 – 3.33 (m, 1H), 3.29 – 3.22 (m, 2H), 2.49 – 2.38 (m, 2H), 2.20 – 2.05 (m, 3H), 1.77 – 1.70 (m, 1H), 1.64 – 1.56 (m, 2H), 1.49 – 1.44 (m, 2H), 1.36 – 1.28 (m, 4H), 1.09 – 1.07 (m, 3H).

**^13^C NMR** (126 MHz, CD_3_OD) δ 176.28, 176.23, 173.16, 172.58, 172.29, 166.09, 145.78, 144.19, 142.48, 141.45, 136.96, 135.57, 133.51, 132.83, 132.67, 132.24, 130.99, 130.65, 130.55, 129.89, 129.75, 128.83, 126.06, 71.78, 71.59, 71.53, 71.51, 71.45, 71.25, 70.59, 70.56, 70.52, 57.38, 52.70, 49.88, 40.36, 40.29, 36.92, 36.80, 35.80, 35.39, 30.75, 30.20, 27.16, 26.83, 15.73.

**HRMS** (ESI+) m/z calcd for C_46_H_61_N_10_O_13_S^+^ [M+H]^+^: 993.4135, found 993.4123.

Synthesis of iodoacetamide-PEG-desthiobiotin (IA-DTB) and chloroacetamide-PEG-desthiobiotin (CA-DTB)

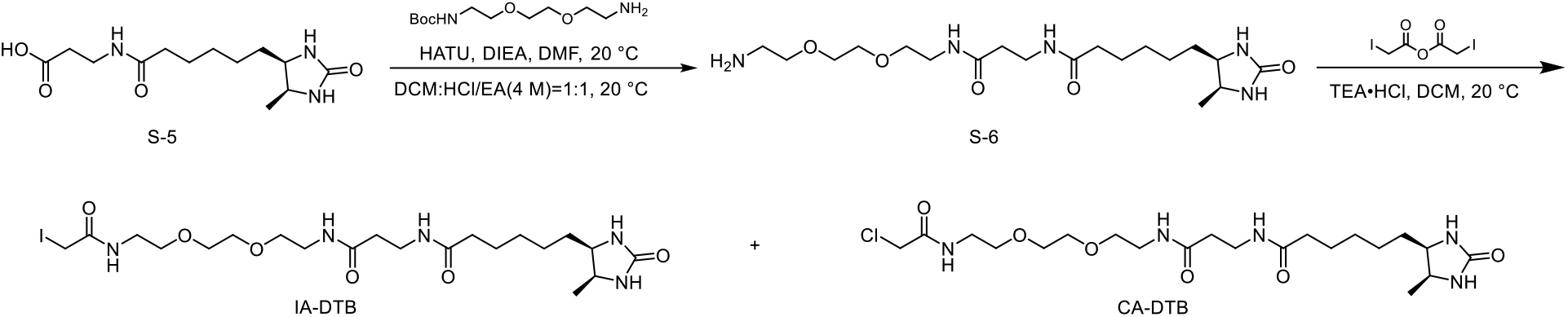

S-5, synthesized by previously reported method^54^, was coupled with N-Boc-2,2’- (ethylenedioxy)diethylamine which followed by deprotection of tert-Butyloxycarbonyl to give S-6. In presence of iodoacetic anhydride and triethylamine hydrochloride, S-6 was stirred in DCM at room temperature for 2 days and finally gave mixture of IA-DTB (10.2 mg, 0.017 mmol, 51%) and CA-DTB (6.6 mg, 0.013 mmol, 33%).

IA-DTB: **^1^H NMR** (500 MHz, CDCl_3_) δ 7.43 (s, 1H), 7.02 (s, 1H), 6.60 (s, 1H), 3.89 – 3.84 (m, 1H), 3.77 – 3.70 (m, 3H), 3.63 – 3.47 (m, 14H), 2.50 (t, *J* = 4.0 Hz, 2H), 2.21 (t, *J* = 8.0 Hz, 2H), 1.68 – 1.65 (m, 2H), 1.46 – 1.32 (m, 6H), 1.15 (d, *J* = 8.0 Hz, 3H).

**HRMS** (ESI+) m/z calcd for C_21_H_39_IN_5_O_6_ ^+^ [M+H]^+^: 584.1940, found 584.1958. CA-DTB: **^1^H NMR** (500 MHz, CDCl_3_) δ 7.19 (s, 1H), 6.79 (s, 1H), 6.37 (s, 1H), 4.08 (s, 2H), 3.88 – 3.84 (m, 1H), 3.73 – 3.70 (m, 1H), 3.64 – 3.60 (m, 6H), 3.58 – 3.52 (m, 6H), 3.48 – 3.44 (m, 2H), 2.46 (t, *J* = 4.0 Hz, 2H), 2.18 (t, *J* = 8.0 Hz, 2H), 1.67 – 1.64 (m, 2H), 1.48 – 1.26 (m, 6H), 1.15 (d, *J* = 4.0 Hz, 3H).

**HRMS** (ESI+) m/z calcd for C_21_H_39_ClN_5_O_6_ ^+^ [M+H]^+^: 492.2583, found 492.2597. Synthesis of iodoacetamide-carboxylate-PEG-desthiobiotin (IA-DTB-COOH)

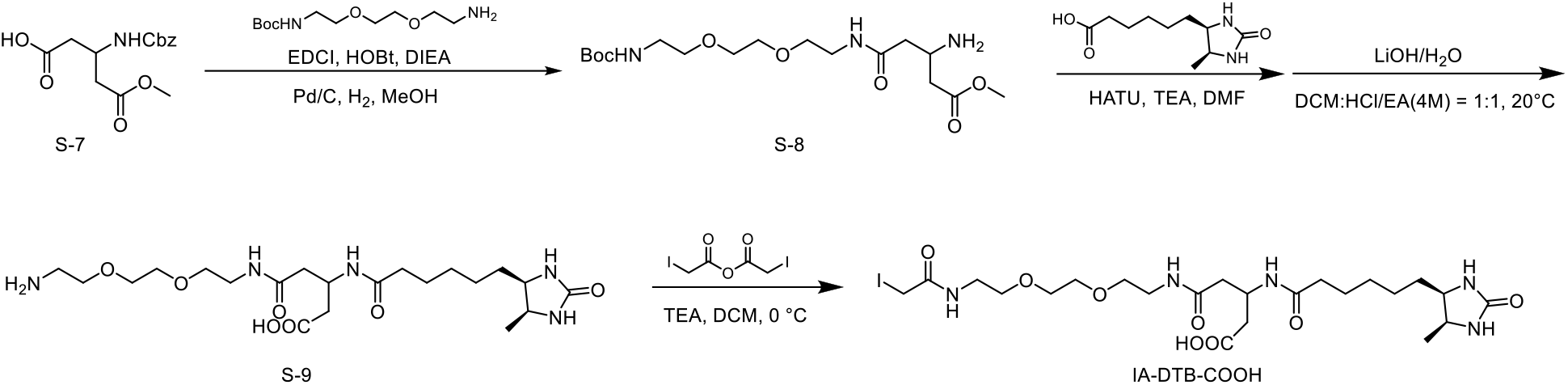

S-7, synthesized by previously reported method^55^, was condensation with N-Boc-2,2’- (ethylenedioxy)diethylamine which then underwent deprotection of N-carbobenzyloxy group to give S-8. Intermediate S-8 went through sequentially coupling with *D*-Desthiobiotin and two steps of hydrolysis provided S-9. By using similar procedure with IA-DTB, IA-DTB-COOH was purified by Pre-HPLC (12.7 mg, 0.020 mmol, 87%).

**^1^H NMR** (500 MHz, CDCl_3_) δ 7.70(s, 1H), 7.22 – 7.15 (m, 1H), 6.00 – 5.84 (s, 1H), 5.32 – 5.09 (m, 1H), 4.63 – 4.39 (m, 1H), 3.66 – 3.26 (m, 12H), 2.44 – 2.35 (m, 2H), 2.13 – 2.02 (m, 2H), 1.47 – 1.23 (m, 11H), 0.96 (d, *J* = 8.0 Hz, 3H).

**HRMS** (ESI+) m/z calcd for C_23_H_41_IN_5_O_8_ ^+^ [M+H]^+^: 642.1994, found 642. 1975. Synthesis of desthiobiotin iodoacetamide (DBIA)

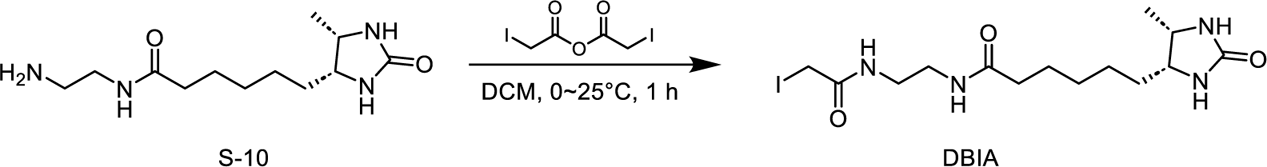

S-10, synthesized by previously reported method^56^, was reacted with iodoacetic anhydride in DCM to afford DBIA (50.0 mg, 0.119 mmol, 63%).

**^1^H NMR** (500 MHz, CDCl_3_) δ 8.26 (s, 1H), 7.78 (s, 1H), 6.29 (s, 1H), 6.11 (s, 1H), 3.63 – 3.57 (m, 3H), 3.49 – 3.46 (m, 1H), 3.07 (s, 4H), 2.04 (t, *J* = 8.0 Hz, 2H), 1.51 – 1.44 (m, 2H), 1.35 – 1.28 (m, 6H), 1.27 – 1.13 (m, 2H), 0.96 (d, *J* = 4.0 Hz, 3H).

**HRMS** (ESI+) m/z calcd for C_14_H_26_IN_4_O_3_ ^+^ [M+H]^+^: 425.1044, found 425.1029.

